# Ventral hippocampus-lateral septum circuitry promotes foraging-related memory

**DOI:** 10.1101/2020.06.16.155721

**Authors:** Léa Décarie-Spain, Clarissa M. Liu, Logan Tierno Lauer, Keshav Subramanian, Alexander G. Bashaw, Molly E. Klug, Isabella H. Gianatiempo, Andrea N. Suarez, Emily E. Noble, Kristen N. Donohue, Alyssa M. Cortella, Joel D. Hahn, Elizabeth A. Davis, Scott E. Kanoski

## Abstract

Remembering the location of a food or water source is essential for survival. Here we demonstrate that spatial memory for food location is reflected in ventral hippocampus (HPCv) neuron activity and is impaired by HPCv lesion. HPCv mediation of foraging-related memory involves downstream lateral septum (LS) signaling, as both reversible and chronic disconnection of HPCv (field CA1) to LS signaling impairs spatial memory retention for the location of either food or water reinforcement. This neural pathway selectively encodes appetitive spatial memory, as HPCv-LS disconnection does not affect aversive reinforcement-based spatial memory in an escape task using the same apparatus. The selectivity of this pathway in promoting foraging-related memory is further supported by results showing that CA1v-LS disconnection does not affect food intake, motivated operant responses for food, anxiety-like behavior, locomotor activity, or social and olfactory-based appetitive learning. Fluorescent in situ hybridization reveals that LS neurons recruited during the appetitive spatial memory task are primarily GABAergic, and multisynaptic anterograde neural pathway tracing and immediate early gene mapping identify the lateral hypothalamic area as a functional downstream target of this pathway. Collective results reveal a novel neural circuit through which the hippocampus selectively mediates memory for the location of appetitive (food or water) but not aversive (escape) reinforcement.

## INTRODUCTION

A survival advantage common to early humans and lower-order mammals is the ability to accurately remember the location of a food or water source in the environment, and efficiently navigate back to a shelter or other safe location. The neurobiological substrates that regulate visuospatial navigation are therefore critical to effectively implement food-directed or water-directed foraging behavior. In rodents, a number of maze tasks have been developed that have been instrumental in identifying brain structures within the telencephalon that are essential for visuospatial mapping and egocentric orientation in the environment, including the hippocampus and the medial entorhinal cortex, respectively (Morris, 1997; Rowland et al., 2016). However, despite that reliably locating food and water sources in the environment is a key selection pressure driving the evolution of visuospatial navigation, the overwhelming majority of rodent model research on the neural substrates of spatial learning and memory have utilized procedures such as the Morris water maze and the Barnes maze that each involve escaping aversive stimulus conditions (Barnes, 1979; Morris, 1984). Furthermore, while single unit recordings have been used to identify specific populations of neurons that subserve distinct navigational functions (e.g., hippocampal “place cells”, medial entorhinal “grid cells”) (Hafting et al., 2005; O’Keefe and Dostrovsky, 1971), and these neurons have been shown to respond to taste (Herzog et al., 2019, 2020; Ho et al., 2011), the bulk of this research has recorded neural activity under neutral conditions void of either appetitive or aversive reinforcement. Very little research has been dedicated to identifying brain regions and neural circuits that may specifically promote spatial memory based on the location of appetitive reinforcers (e.g., food, water), as well as the extent to which the nature of the reinforcement is a critical factor in deciphering the brain’s control of visuospatial navigation.

Rodent model research investigating brain regions that mediate visuospatial navigational memory has predominantly focused on the anterior and “dorsal” subregion of the hippocampus (septal pole; HPCd). However, the posterior and “ventral” hippocampus subregion (temporal pole; HPCv), while classically associated with stress- and affect-associated memory processes (Fanselow and Dong, 2010), also plays a role in visuospatial learning and memory (de Hoz et al., 2003; Keinath et al., 2014; Kjelstrup et al., 2008). For example, under some testing conditions, selective HPCv lesions impair spatial memory performance in the Morris water maze (Ferbinteanu et al., 2003; de Hoz et al., 2003). Moreover, place cells that are responsive to location changes in the visuospatial environment are present within both the HPCd and HPCv pyramidal layers, with a linear increase in the spatial scale of representation from the dorsal to the ventral pole (Kjelstrup et al., 2008). Despite a common role for the HPCd and HPCv in mediating spatial memory, there is also evidence for a functional distinction between the subregions (Fanselow and Dong, 2010; Kanoski and Grill, 2017; Moser and Moser, 1998). For instance, lesions of the HPCv but not the HPCd alter stress responses (Henke, 1990) and anxiety-like behavior (Kjelstrup et al., 2002), whereas HPCd but not HPCv lesions impair spatial memory in an incidental (nonreinforced) procedure (Gaskin et al., 2009). These two HPC subregions also have disparate neuroanatomical connectivity, supporting a framework for a functional diversity in which the HPCd preferentially processes cortical-derived sensory information and the HPCv preferentially processes metabolic and limbic-derived affective information (Fanselow and Dong, 2010; Kanoski and Grill, 2017). These functional distinctions are further supported by the generation of the hippocampus gene expression atlas, which provides a comprehensive integration of gene expression and connectivity across the HPC septo-temporal axis (Bienkowski et al., 2018; Gergues et al., 2020). Given that both the HPCv and the HPCd participate in spatial memory but have distinct neuroanatomical connectivity and contributions when it comes to regulating other cognitive and mnemonic processes, it is feasible that the HPCv and HPCd support different forms of spatial memory depending on the type of reinforcement and/or the context associated with the behavior.

Recent findings identify the HPCv as a critical brain substrate in regulating feeding behavior and food-directed memory processes. Reversible inactivation of HPCv neurons after a meal increases the size of and reduces the latency to consume a subsequent meal (Hannapel et al., 2019, 2017). In addition, receptors for feeding-related hormones are more abundantly expressed in the HPCv compared to the HPCd (e.g., ghrelin receptors (Mani et al., 2014; Zigman et al., 2006), glucagon-like peptide-1 [GLP-1] receptors (Merchenthaler et al., 1999)), and these HPCv endocrine and neuropeptide receptor systems alter food intake and feeding-related memory (Hsu et al., 2015a, 2018a, 2018b; Kanoski et al., 2011, 2013; Suarez et al., 2020). Olfactory information, which is intimately connected with feeding behavior, is also preferentially processed within the HPCv compared with the HPCd (Aqrabawi and Kim, 2018; Aqrabawi et al., 2016). The HPCv field CA1 (CA1v), specifically, is bidirectionally connected to brain regions that process olfactory information (Fanselow and Dong, 2010; De La Rosa-Prieto et al., 2009; Petrovich et al., 2001), and CA1v neurons respond more robustly to olfactory contextual cues compared with CA1d (Keinath et al., 2014). Given these appetite-relevant HPCv neuroanatomical connections, endocrine and neuropeptide receptor expression profiles, and functional evidence linking this subregion with food-motivated behaviors, we hypothesize that HPCv mediation of visuospatial memory is preferentially biased to appetitive food and water-reinforced foraging behavior.

HPCv pyramidal neurons have extensive projection targets throughout the brain (Swanson and Cowan, 1977), yet the functional relevance of these pathways is poorly understood. The lateral septum is a robust target of HPCv glutamatergic pyramidal neurons (Arszovszki et al., 2014; Risold and Swanson, 1996). Given that this pathway has been shown to affect feeding behavior (Kosugi et al., 2021; Sweeney and Yang, 2015) and that neuroplastic changes occur in the LS after learning a spatial memory task (Garcia et al., 1993; Jaffard et al., 1996), we predict that that HPCv to LS projections participate in regulating foraging-related spatial memory for food and water location. To investigate this hypothesis, we developed two appetitive reinforcement-based spatial foraging behavioral tasks that allow for direct comparison with an aversive reinforcement-based task of similar difficulty that uses the same apparatus and spatial cues. First, foraging-related spatial memory was assessed following neurotoxic ablation of the HPCv, and CA1v bulk intracellular calcium activity was recorded during memory probe performance in intact control animals. Performance in these tasks was next assessed following pathway-specific dual viral-based reversible (chemogenetic inhibition) or chronic (targeted apoptosis) disconnection of the CA1v to LS pathway. To further expand neural network knowledge on CA1v to LS signaling, we used conditional viral-based neuroanatomical tracing strategies to identify both first-order collateral and second-order targets of LS-projecting CA1v neurons, as well as immediate early gene mapping approaches to confirm the functional relevance of downstream pathways. Collective results from the present study identify novel neural circuitry of specific relevance to foraging behavior.

## RESULTS

### Ventral hippocampus (HPCv) neurons encode memory for the spatial location of food

To examine the importance of the HPCv in memory for the spatial location of food reinforcement, HPCv bulk calcium-dependent activity was recorded in animals performing a food-reinforced appetitive visuospatial memory task that utilizes a Barnes maze apparatus (diagram in Fig. 1A). Fiber photometry recordings in the HPCv (representative injection site in Fig. 1B-C) revealed HPCv activity fluctuations in response to escape hole investigations during the appetitive visuospatial memory task memory probe in which no food was present (Fig. 1D; time x investigation type p<0.05). HPCv changes in neuron activity immediately following escape hole investigation differ between investigation type (correct vs. incorrect; Fig. 1E; p<0.05, cohen’s d=-1.0582), such that 5-second calcium activity increased following correct hole investigations yet decreased following incorrect hole investigations. In addition, a significant correlation was found between changes in HPCv activity 5s following a hole investigation and distance from the appropriate escape hole (Fig. 1F; p<0.01, r^2^=0.08584). To verify whether the HPCv is necessary for food location-based spatial memory, another group of animals received bilateral *N*-Methyl-D-aspartate excitotoxic lesions of the HPCv or bilateral sham injections (histological analyses for the neuron-specific NeuN antigen in Fig. 1G) and were tested the appetitive visuospatial memory task (representative behavioral traces in Fig. 1H). Results revealed no significant differences in errors (incorrect hole investigations) (Fig. 1I) or latency to locate the food source hole (Fig. 1J) during training. However, memory probe results show that animals with HPCv lesions decreased the ratio of correct + adjacent / total holes explored compared with controls (Fig. 1K; Control vs. HPCv lesion, p<0.05, cohen’s d=-1.0174, t-test; Control group different from chance performance, p<0.001, one-sample t-test) indicating impaired memory for the spatial location of food reinforcement.

**Figure 1.**
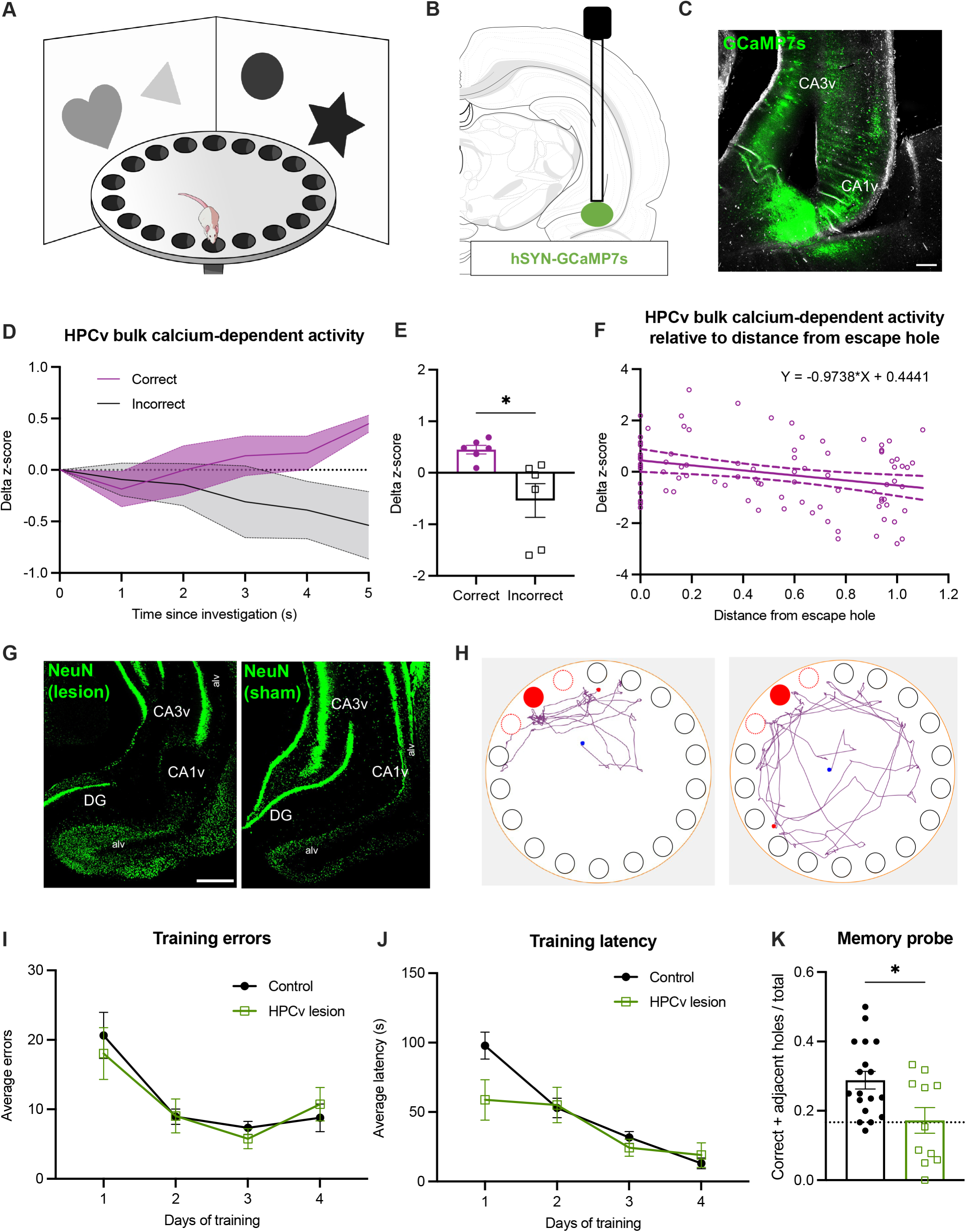
Bilateral HPCv lesions impair spatial memory for food location. Spatial foraging task apparatus (A). Diagram of stereotaxic viral infusion (AAV-hSYN-GCaMP7s) and placement of optic fiber in the HPCv for bulk photometry recordings (B). Representative histology of GCaMP7s expression in HPCv neurons (C; scale bars 100 μm). Variation in HPCv bulk calcium-dependent activity following a correct or incorrect investigation during a spatial food seeking task (Time x Investigation type p<0.05; D). Delta z-score 5s following a correct versus incorrect investigation during a spatial food seeking task (p<0.05; E). Simple linear regression of delta z-score 5s following investigation relative to distance of investigated hole from correct escape hole (p<0.01; F). Representative HPCv lesion histology with NeuN immunohistochemistry (G; scale bars 500μm). Representative navigation paths of a control animal preferentially investigating correct (filled red) and adjacent holes (outlined orange) during spatial foraging memory probe (H, left). Representative navigation path of HPCv lesioned animal during spatial foraging memory probe (H, right). Bilateral HPCv lesions did not impair learning of the spatial foraging task compared to controls, as measured by errors before locating correct hole during task training (I) and latency to locate correct hole during task training (J). Bilateral HPCv lesions impaired retention of the spatial foraging task, as measured by the ratio of investigation of correct plus adjacent holes over total investigated during the first minute of the task (p<0.05; K). Dotted line indicates chance performance level (0.167). For graphs 1D-F, n=6 (within-subjects). For graphs 1I-K, lesion n=11, control n=18. All values expressed as mean +/- SEM.

Post-surgical analyses of food intake (Supp. Fig. 1A) and body weight (Supp. Fig. 1B) found no significant group differences in these measures over time. These collective results indicate that the HPCv is required for memory retention but not learning of the spatial location of food in the environment, and that HPCv neural activity is modulated by appetitive spatial memory accuracy.

### Ventral hippocampus CA1 (CA1v) projections to the lateral septum (LS) mediate appetitive spatial memory for food or water location, but not aversive spatial memory for escape location

To identify a specific neural pathway mediating the results described above, we investigated the functional relevance of the CA1v to LS signaling in learning and recalling the spatial location of either food or water reinforcement using conditional dual viral approaches to either reversibly (via cre-dependent pathway-specific inhibitory chemogenetics with CNO administration prior to the memory probe; diagram of approach in Fig. 2A) or permanently (via cre-dependent pathway-specific caspase-induced lesions; diagram of approach in Fig. 2B) disconnect CA1v to LS communication. Results from the appetitive food seeking spatial memory task reveal no significant group differences during training in either errors before locating the correct hole (Fig. 2D) or latency to locate the food source hole (Fig. 2E). However, memory probe results demonstrate that both acute (pathway-specific DREADDs) and chronic (pathway-specific caspase) CA1v to LS disconnection decreased the ratio of correct + adjacent / total holes explored compared with controls (Fig. 2F; Control vs. DREADDs and Control vs. Caspase, p<0.05, cohen’s d=-1.142; Control different from chance, p<0.0001, Caspase different from chance, p<0.05, One-sample t-test), indicating impaired memory for the spatial location of food reinforcement.

**Figure 2.**
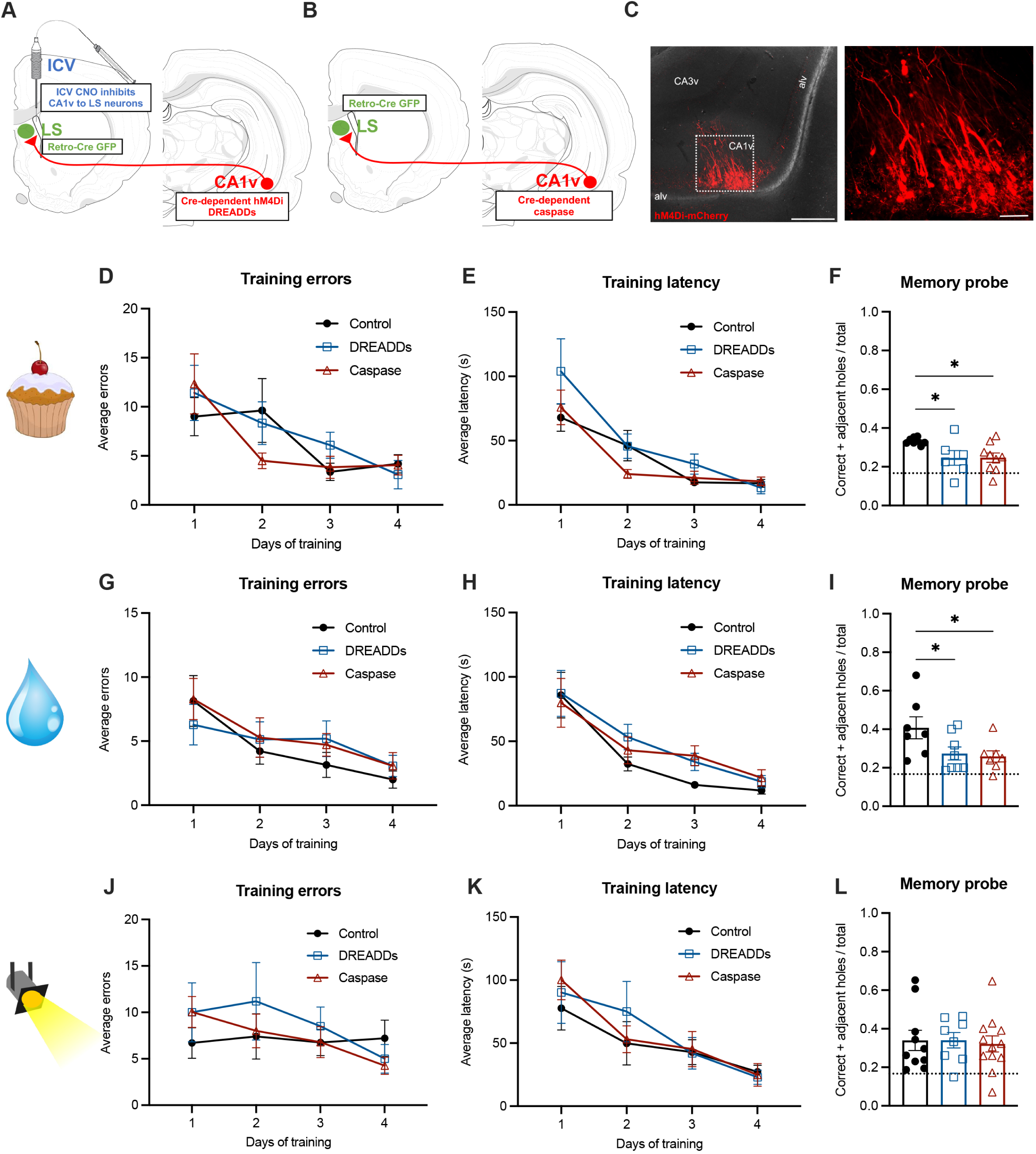
Reversible and chronic CA1v to LS neural disconnection impairs spatial memory for food and water location but not for escape location. Diagram of dual viral approach using a cre-dependent inhibitory DREADDs (AAV-hSyn-DIO-hM4D(Gi)-mCherry) approach to reversibly disconnect CA1v to LS neural pathway (with CNO injected 1hr before memory probe) (A). Diagram of dual viral approach using a cre-dependent caspase (AAV1-Flex-taCasp3-TEVp) approach to chronically disconnect CA1v to LS neural pathway (B). Representative injection site in CA1v demonstrating LS-projecting neurons infected with inhibitory DREADDs, which simultaneously drives expression of a fluorescent mCherry transgene (C; scale bars 500μm [left] and 100μm [right]). Neither reversible (DREADDs) nor chronic (caspase) disconnection of the CA1v to LS pathway impaired learning of the spatial food seeking task compared to controls, as measured by errors before correct hole during task training (D) and latency to correct hole during task training (E). Both reversible and chronic disconnection of the CA1v to LS pathway impaired retention of the food location as measured by the ratio of investigation of correct plus adjacent holes over total investigations during entire two minutes of the task (p<0.05; F). Likewise, reversible (DREADDs) and chronic (caspase) disconnection of the CA1v to LS pathway did not impair learning of the spatial water seeking task compared to controls, as measured by errors before correct hole during task training (G) and latency to correct hole during task training (H), but impaired memory retention of the water location during the probe (I). In contrast, disconnection of the CA1v to LS pathway either reversibly (DREADDs) or chronically (caspase) did not impair performance on the spatial escape task. There were no differences in learning as measured by errors before correct hole during task training (J) and latency to correct hole during task training (K). Unlike the spatial foraging task, retention of the spatial escape task was not impaired by reversible nor chronic disconnection of the CA1v to LS pathway (L). For graphs 2E-G (CA1v to LS disconnect cohort 1), DREADDs n=6, caspase n=10, control n=8. For graphs 2H-J (CA1v to LS disconnect cohort 3), DREADDs n=8, caspase n=8, control n=7). For graphs 2K-M (CA1v to LS disconnect cohort 2), DREADDs n=8, caspase n=12, control n=10. Dotted line indicates chance performance level (0.167). All values expressed as mean +/− SEM.

Results from an analogous appetitive water seeking spatial memory task in a separate cohort of water-restricted rats also reveal no significant group differences during training in either errors before locating the correct hole (Fig. 2G) or latency to locate the water source hole (Fig. 2H). However, a significant impairment was observed following either acute or chronic CA1v to LS disconnection of the ratio of correct + adjacent / total holes explored compared with controls (Fig 2I; Control vs. DREADDs and Control vs. Caspase, p<0.05, cohen’s d=-1.18; Control different from chance, p<0.01, DREADDs and Caspase different from chance, p<0.05, One-sample t-test), indicating impaired memory for the spatial location of water reinforcement.

Histological analyses confirmed successful viral transfection of LS-projecting CA1v neurons with DREADDs (Fig. 2C), as well as successful lesioning of LS-projecting CA1v neurons for the caspase approach (Supplemental Fig. 2G-H). The reversible DREADDs-based disconnection approach was further confirmed by c-Fos analyses in the LS, where the number of c-Fos immunoreactive neurons in the LS in response to the food reinforcement-based spatial memory probe test was reduced following CNO treatment relative to vehicle/control treatment (Supp. Fig. 2A-C; p<0.01, cohen’s d=-3.9578). We further confirmed that ICV administration of CNO failed to influence LS neuronal activation in the absence of inhibitory DREADDs (Supp. Fig. 2D-F). Post-surgical analyses of food intake (Supp. Fig 1C) and body weight (Supp. Fig 1D) found no group differences in these measures over time (in the absence of CNO treatment). These data demonstrate that the CA1v-LS disconnection approaches were successful, that CA1v to LS communication is critical for remembering the spatial location of food and water, and that these effects are unlikely to be based on differences in energy status or food motivation.

To evaluate whether CA1v to LS signaling is specifically involved in appetitive reinforcement-based spatial memory (food or water) vs. spatial memory in general, in a separate cohort of animals we tested the effect of reversible and chronic CA1v to LS disconnection in a spatial learning task based on aversive reinforcement rather than food or water. Importantly, this task uses the same apparatus and visuospatial cues as the spatial location food and water memory tasks described above, but the animals are motivated to locate the tunnel to escape mildly aversive stimuli (bright lights, loud noise) with no food or water reinforcement in the tunnel. Training results revealed no significant group differences in errors before correct hole (Fig. 2J) nor latency to locate the escape hole (Fig. 2K). During the memory probe test, there were no group differences in the ratio of correct + adjacent / total holes investigated (Fig. 2L; Control, DREADDs and Caspase different from chance, p<0.01, One-sample t-test). Post-surgical analyses of food intake (Supp. Fig. 1E) and body weight (Supp. Fig. 1F) found no group differences in these measures over time. These collective findings suggest that CA1v to LS signaling specifically mediates spatial memory in a reinforcement-dependent manner, playing a role in appetitive (food or water reinforcement) but not aversive (escape-based) spatial memory.

### Disconnection of CA1v to LS signaling does not influence food intake, food-motivated operant responding, nonspatial HPC-dependent appetitive memory, anxiety-like behavior, or locomotor activity

While the results above suggest that the impairments in food or water-based spatial memory were not based on long-term changes in food intake or body weight, these experiments were conducted under conditions of either chronic food or water restriction. To examine the role of CA1v to LS signaling on feeding behavior under free-feeding conditions, we examined meal pattern feeding analyses in the caspase vs. the control group (chronic CA1v-LS disconnection), and in the DREADDs group following CNO or vehicle (within-subjects design; reversible CA1v-LS disconnection). Results averaged over 5d revealed no significant effect of chronic CA1v to LS disconnection (caspase group) on 24h meal frequency, meal size, meal duration, or 24h food intake in comparison to controls (Supp. Figs. 3A-D). Similarly, CNO-induced acute reversible silencing of LS-projecting CA1v neurons (DREADDs group) did not significantly affect 24h meal frequency, meal size, meal duration or 24h food intake relative to vehicle treatment (Supp. Fig. 3E-H). Further, CNO-induced acute disconnection of the CA1v to LS pathway had no impact on effort-based lever press responding for sucrose rewards in a progressive ratio reinforcement procedure (Supp. Fig. 3I). Collectively these findings show that neither reversible nor chronic CA1v to LS disconnection influences food intake under free-feeding conditions nor food motivation in a non-visuospatial task.

The HPCv plays an important role in the social transmission of food preference procedure (STFP) (Carballo-Márquez et al., 2009; Countryman et al., 2005; Hsu et al., 2018c), which tests socially-mediated food-related memory based on previous exposure to socially-communicated olfactory cues (diagram in Supp. Fig. 4A). However, results revealed that neither chronic nor acute disconnection of CA1v to LS signaling affected the STFP preference ratio between groups (Supp. Fig. 4B), with all three groups performing significantly above chance (based on within-group one sample t-tests, p<0.05).

HPCv to LS circuitry has been shown to play a role in mediating anxiety and stress behavior (Parfitt et al., 2017; Trent and Menard, 2010). Given that altered anxiety-like behavior and/or locomotion could produce behavioral changes that may lead to an apparent spatial memory deficit in any of the spatial learning tasks used, we examined whether CA1v to LS disconnection yields differences in anxiety-like behavior (zero maze task; diagram of apparatus in Supp. Fig. 4C) and/or locomotion (open field test). Zero maze results showed no significant group differences in time spent in the open zones (Supp. Fig. 4D) nor in the number of open zone entries (Supp. Fig. 4E). In addition, CA1v to LS disconnection had no effect on general locomotor activity in the open field test (Supp. Fig. 4F). These collective results suggest that the observed deficits in food- and water-seeking spatial memory are not secondary to impaired nonspatial appetitive learning, general differences in anxiety-like behavior, or locomotor impairments.

### Collateral medial prefrontal cortex projections from LS-projecting CA1v neurons do not contribute to appetitive spatial memory

In addition to the LS, CA1v neurons also robustly project to the medial prefrontal cortex (mPFC), a pathway that we have previously shown to be involved in feeding behavior (Hsu et al., 2018a). To examine whether LS-projecting CA1v neurons have collateral targets in the mPFC, or in other brain regions, we utilized a conditional dual viral neural pathway tracing approach that identifies collateral targets of a specific monosynaptic pathway (CA1v->LS; diagram of approach in Fig. 3A representative CA1v and LS injection sites in Fig. 3B [left and middle]). Results revealed that LS-projecting CA1v neurons also project to the mPFC (Fig. 3B (right)), whereas very minimal labeling was observed in other brain regions throughout the neuraxis. Thus, it may be the case that the impaired spatial memory for food location observed following either reversible or chronic CA1v to LS disconnection are based, in part, on CA1v to mPFC signaling from the same LS-projecting CA1v neurons.

**Figure 3.**
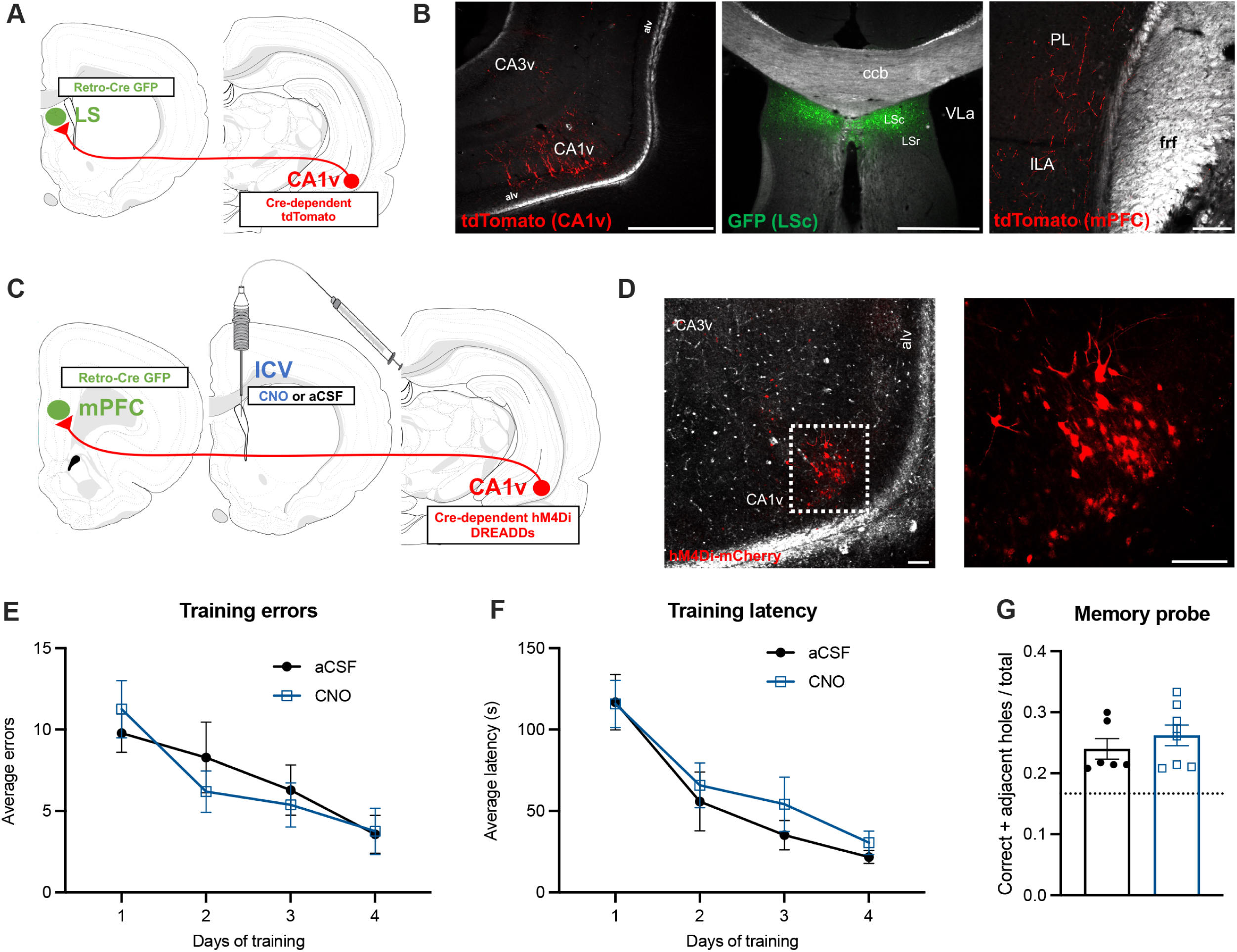
CA1v to mPFC projections do not contribute to appetitive spatial memory. Representative CA1v injection site from collateral identification approach (A, left; scale bar 500μm). Representative LS injection site from collateral identification approach in relation to the caudal (LSc) and rostral (LSr) subregions, the corpus callosum body (ccb) and lateral ventricle (VLa) (A, middle; scale bar 500μm). Representative image collateral axons of the CA1v to LS pathway located in the prelimbic (PL) and infralimbic (ILA) areas of the mPFC (A, right; scale bar 50μm). Diagram of dual viral approach to identify collateral targets of the CA1v to LS neural pathway (B). Diagram of dual viral approach using cre-dependent inhibitory DREADDs (AAV-hSyn-DIO-hM4D(Gi)-mCherry) to acutely disconnect the CA1v to mPFC neural pathway (with CNO or aCSF injected 1hr before memory probe) (C). Representative injection site in CA1v demonstrating mPFC-projecting neurons infected with inhibitory DREADDs, which simultaneously drives expression of a fluorescent mCherry transgene (D; scale bars 100μm). Disconnection of the CA1v to mPFC neural pathway did not influence learning of the spatial foraging task compared to controls, as measured by errors before correct hole during training (E) and latency to correct hole during training (F). Disconnection of the CA1v to mPFC pathway did not influence retention of the spatial foraging task in the memory probe (G). For graphs 3E-G, aCSF n=6, CNO n=8. All values expressed as mean +/− SEM.

We next sought to test the functional relevance to food-directed spatial memory of a CA1v-originating neural pathway (CA1v to mPFC) that is a collateral target of LS-projecting CA1v neurons. Inhibitory DREADDs were expressed in mPFC-projecting CA1v neurons (diagram of approach in Fig. 3C and representative CA1v injection site in Fig. 3D), and animals received an ICV infusion of CNO or aCSF 1h prior to the memory probe. Training results from the appetitive spatial memory task revealed no group differences in errors before locating the correct hole (Fig. 3E) or latency to locate the food-baited hole (Fig. 3F). Importantly, memory probe results demonstrate that CA1v to mPFC disconnection (via CNO treatment) did not alter the ratio of correct + adjacent / total holes explored (Fig. 3G; aCSF different from chance, p<0.01, CNO different from chance, p<0.001, One-sample t-test) compared with controls (aCSF). These data collectively demonstrate that CA1v to LS mediation of foraging-related memory does not require collateral projections to the mPFC neural pathway and that the impact of ICV CNO administration on memory performance is not generalized to the inhibition of any CA1v projections.

### Appetitive spatial memory engages LS GABAergic neurons and the lateral hypothalamic area (LHA) downstream of HPCv-LS signaling

Fluorescent *in situ* hybridization analyses were used to determine the neurotransmitter phenotype of c-Fos+ LS neurons that were activated during the appetitive food seeking spatial memory probe test. Results revealed that ∼10% of c-Fos+ LS neurons express ChAT (a cholinergic marker; Fig. 4A), ∼88% of c-Fos+ LS neurons express GAD1 (a GABAergic marker; Fig. 4B), and only ∼2% of c-Fos+ LS neurons express VGLUT2 (a glutamatergic marker; Fig. 4C). These results suggest that LS neurons engaged during the spatial memory probe are primarily GABAergic (Fig. 4D).

**Figure 4.**
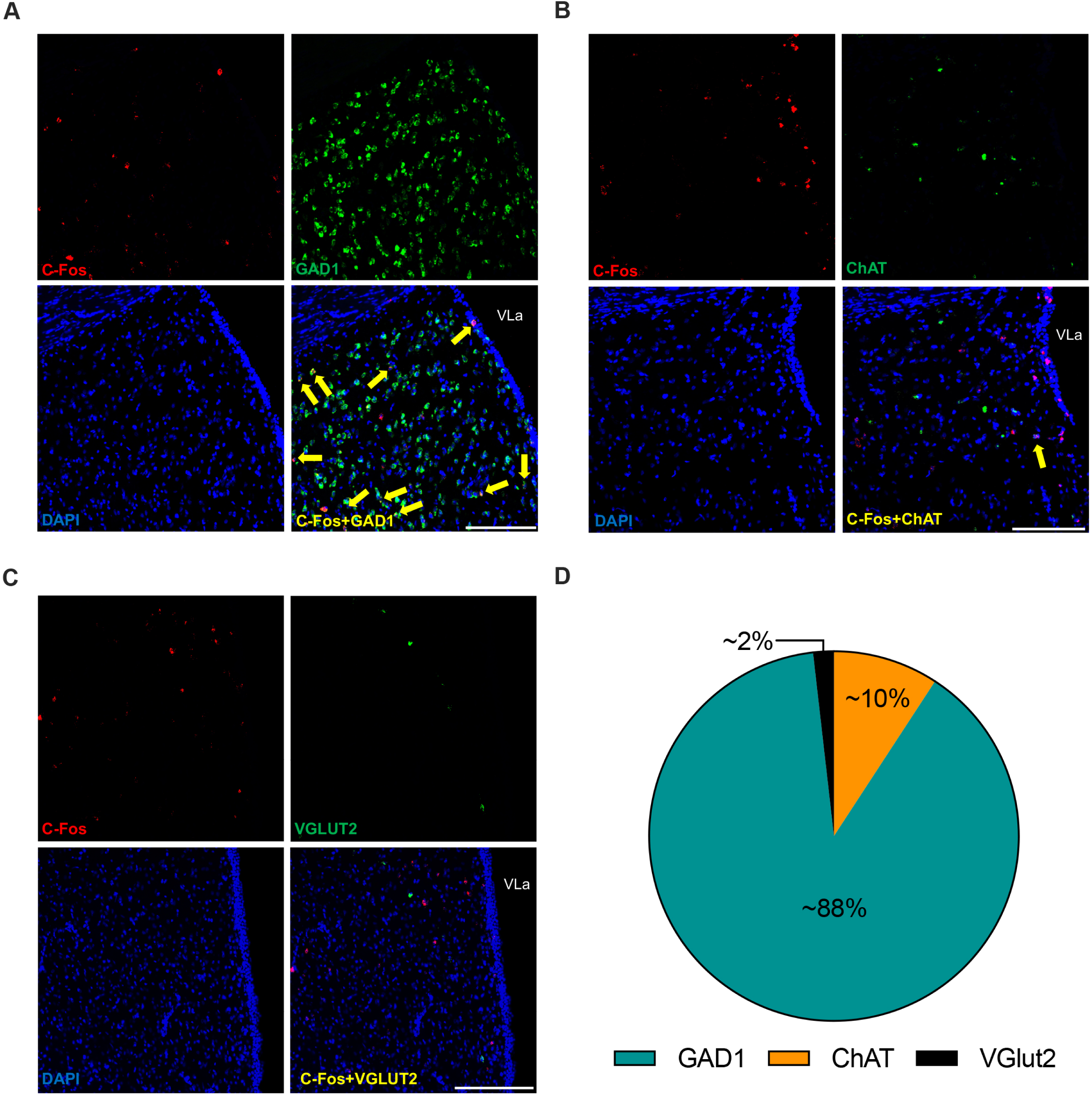
LS neurons activated during food-seeking memory probe are predominantly GABAergic. c-Fos and ChAT colocalization was observed in ∼2% of LS neurons (A; scale bar 500μm), c-Fos and GAD1 colocalization was observed in ∼88% of LS neurons (B; scale bar 500μm), and c-Fos and VGLUT2 colocalization was observed in ∼10% of LS neurons (C; scale bar 500μm) following the spatial memory probe test (for food reinforcement). For graphs A-C, quantification from DREADDs vehicle group, n=3.

A dual viral tracing strategy (diagram of approach in Fig. 5A; representative LS injection site in Fig. 5B) was utilized to identify downstream targets of LS-projecting CA1v neurons. Results reveal that the LHA is the strongest second-order target of the CA1v to LS-projecting neurons (Fig. 5C-D). Quantitative analyses using a custom-built data-entry platform (Axiome C, created by JDH) are summarized graphically for a representative animal on a brain flatmap summary diagram (Fig. 5D) for the hemisphere ipsilateral (top) and contralateral (bottom) to the injection sites (see Supplementary Table 1 for data from complete forebrain analyses). Retrograde transfection of the AAVs from the injection sites was not observed, as tdTomato+ cell bodies were only observed in the LS following complete forebrain histological analyses. Additional confirmation of the LHA as a second-order target was provided via a co-injection (COIN) strategy (Suarez et al., 2018a; Thompson and Swanson, 2010). This approach involved injections of a monosynaptic retrograde tracer, fluorogold (FG), in the LHA (representative injection site in Fig. 5E) and injection of a monosynaptic anterograde tracer, AAV1-GFP, in the CA1v (representative injection site in Fig. 5F), which resulted in the observance of FG-GFP appositions in approximately 57.57% ± 5.04% of FG cell bodies in the LS, thus further confirming this multisynaptic pathway (Fig. 5G).

**Figure 5.**
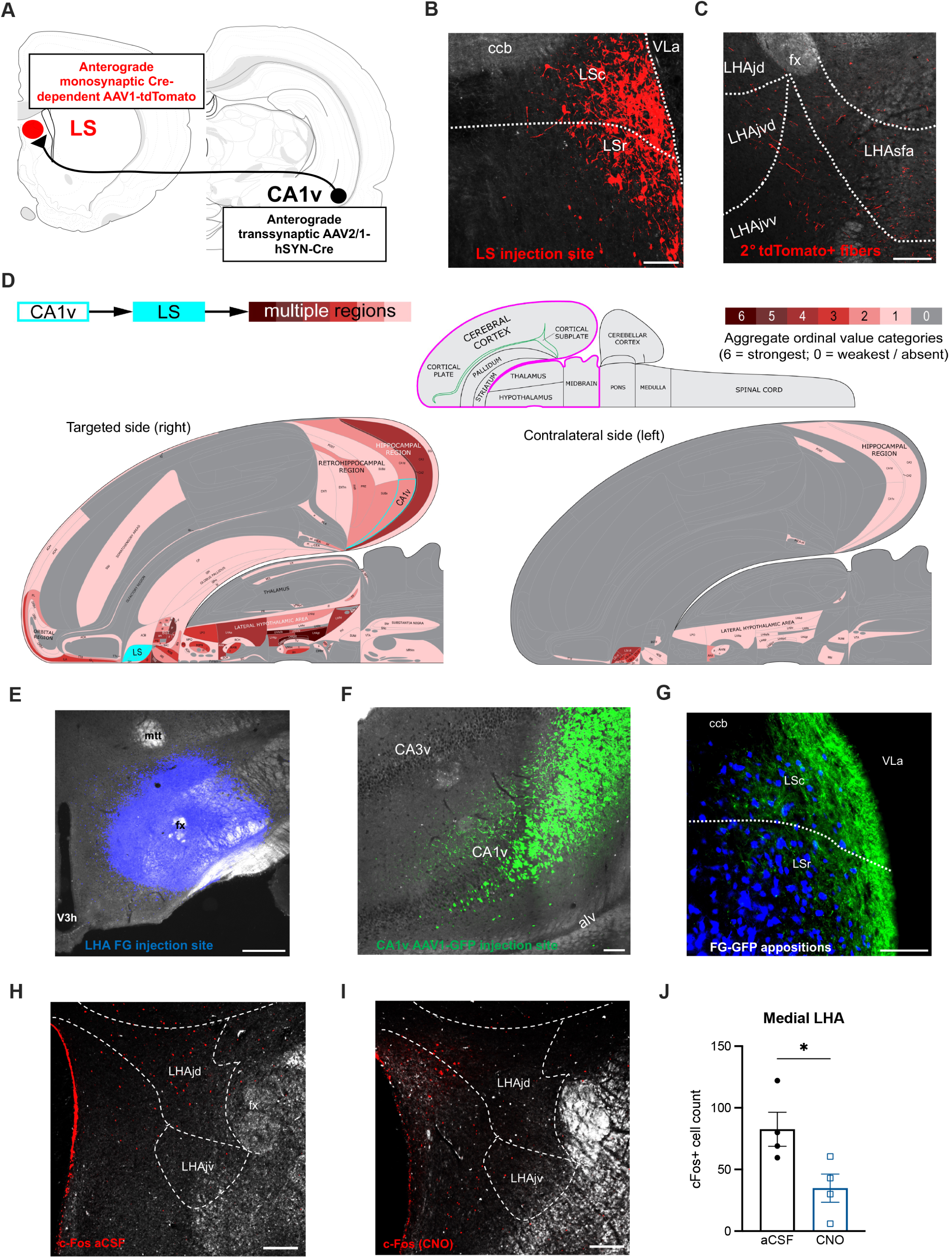
Identification of second-order neural projections downstream of CA1v to LS projections. Diagram of dual viral approach to identify brain regions that are second-order (2°) targets of the CA1v to LS neural pathway (A). Representative LS injection site from second order identification approach in relation to the caudal (LSc) and rostral (LSr) subregions, the corpus callosum body (ccb) and lateral ventricle (VLa) (B, left; scale bar 100μm). Representative image of second-order fibers of the CA1v to LS pathway within the juxtadorsomedial (jd), juxtaventromedial dorsal (jvd), juxtaventromedial ventral and subfornical anterior (sfa) subregions of the LHA (B, right; scale bar 200μm). Summary of the projection targets of LS neurons that receive input from CA1v (D; expanded information in Supplementary Table 1). The outputs of the right side of LS neurons receiving CA1v input are represented at the macroscale (gray matter region resolution) on a partial flatmap representation of the rat forebrain, adapted from (Hahn et al., 2021). Connection weights are represented block colors for each region following an ordinal scale ranging from weakest (0 = very weak or absent) to strongest (6 = strong), as there were no 7 (very strong) values. The inset at lower left represents one side of the brain with the part represented in the upper diagrams outlined in magenta. The COIN approach for further confirmation of second-order tracing strategy is depicted with FG injection site in the LHA (E; scale bar 500μm), representative AAV1-GFP injection site in the CA1v (F; scale bar 100μm), and FG-GFP appositions in the LS (G; scale bar 100μm). Representative histology of c-Fos expression in the medial LHA (LHAjd and LHAjv) following ICV infusion of aCSF (H; scale bar 200μm). Representative histology of c-Fos expression in the medial LHA (LHAjd and LHAjv) following ICV infusion of CNO to inhibit LS-projecting CA1v neurons (I; scale bar 200μm). Counts of c-Fos positive cells in the medial LHA (LHAjd and LHAjv) are reduced with ICV administration of CNO (J; p<0.05). For graph J, aCSF n=4 and CNO n=4. All values expressed as mean +/− SEM.

The functional relevance of this pathway to appetitive spatial memory is provided by immediate early gene mapping of c-Fos+ cells in the LHA subregions that are most densely innervated downstream of CA1v->LS signaling (juxtadorsomedial and juxtaventromedial regions, LHAjd and LHAjv), with results revealing reduced LHA c-Fos expression following appetitive spatial memory probe testing in animals receiving chemogenetic-mediated CA1v-LS disconnection (ICV CNO-injected rats) relative to aCSF-injected controls (Fig. 5H-J; p<0.05, cohen’s d=1.8833). In contrast, medial LHA c-Fos expression following the appetitive spatial memory probe did not differ between by treatment (ICV CNO vs. ICV aCSF) in rats without expression of chemogenetic inhibitory receptors in LS-projecting CA1v neurons (Supp. Fig. 5), thereby further validating the specificity of CNO actions to the inhibitory DREADDS receptor.

Immunohistochemistry analyses were used to determine if acute disconnection of the CA1v-LS pathway influences c-Fos expression of cells co-expressing the orexin or melanin-concentrating hormone (MCH) neuropeptides in the medial LHA, each of which is associated with appetite-promoting effects (Barson et al., 2013; Lord et al., 2021). Results revealed approximately 25% of Orexin+ cells in the medial LHA co-express c-Fos following the appetitive spatial memory probe, and this effect did not differ by aCSF vs. CNO treatment (Supp. Fig. 6). A similar analysis in MCH+ cells of the medial LHA identified only a marginal proportion of MCH+ c-Fos+ neurons (≤3%, data not shown), thus additional analysis was not conducted. These findings suggest that CA1v-LS signaling does not engage downstream orexin or MCH LHA signaling in the context of appetitive foraging-related memory.

## DISCUSSION

Memory for the physical location of a food or water source is adaptive for maintaining adequate energy supply for reproduction and survival. However, the neural circuits mediating this complex behavior are not well understood as research on visuospatial memory has predominantly used tasks with either aversive or neutral/passive reinforcement. Moreover, whether the neural circuitry mediating spatial memory function is divergent based on the nature (e.g., appetitive vs. aversive) or modality (e.g., food vs. water) of the reinforcement has not been systematically investigated. The present study illustrated the critical role the HPCv plays in an appetitive spatial memory task for food location using both bulk regional calcium-dependent activity recordings and a lesion approach. A monosynaptic CA1v to LS pathway is further identified as a necessary substrate for food- and water-motivated spatial memory. Moreover, the selectivity of the CA1v-LS pathway in promoting spatial memory for food and water location is supported by results showing that disconnection of this pathway did not affect performance in an escape-motivated spatial memory task of comparable difficulty conducted in the same apparatus, feeding behavior, anxiety-like behavior, locomotor activity, motivated operant responding for palatable food, or olfactory and social-based appetitive memory. Fluorescent in situ hybridization analyses also demonstrate that LS neurons engaged during appetitive spatial memory testing are primarily GABAergic. Viral-based pathway tracing revealed that LS-projecting CA1v neurons also send collateral projections to the mPFC, however, functional disconnection of CA1v-mPFC signaling did not impair spatial memory for food location. Utilization of an anterograde multisynaptic viral tracing approach and quantitative forebrain-wide axon mapping analyses revealed that the CA1v-LS pathway sends second-order projections to various feeding and reward-relevant neural substrates, particularly the LHA, and immediate early gene functional mapping analyses confirm that this multisynaptic pathway is relevant for appetitive spatial memory. Collectively, these data establish monosynaptic CA1v to LS signaling as an essential pathway for spatial memory-based foraging behavior, and we further identify a neural network of interconnected feeding-relevant substrates that are collateral and/or downstream targets of this pathway.

Historically, the hippocampus has been divided into the dorsal and ventral subregions that have both distinct and overlapping functions, anatomical connections, and genetic and receptor expression profiles (Bienkowski et al., 2018; Fanselow and Dong, 2010; Kanoski and Grill, 2017; Moser and Moser, 1998; Swanson and Cowan, 1977). The HPCd has been predominantly implicated in facilitating spatial memory, whereas the HPCv is linked more with stress responses, affective behavior, and energy balance. However, a recent study found that both the HPCd and HPCv were critical for food reward-directed spatial navigation in an obstacle-rich complex environment (Contreras et al., 2018), and under certain testing conditions, spatial learning and memory in an aversive reinforcement-based water maze also requires both subregions of the HPC (Lee et al., 2019). Here we expand this literature by identifying a specific HPCv-originating pathway (CA1v->LS) that is selectively involved in spatial memory for the location of appetitive (food or water) reinforcement. These novel findings suggest that the modality of the reinforcement should be considered when investigating the longitudinally organized processing of spatial information across the dorsal-ventral HPC axis.

While HPCv neurons project to multiple regions throughout the neuraxis, projections to the LS are particularly robust (Risold and Swanson, 1996). The LS has been implicated in food anticipatory/seeking behavior, where metabolic activation of the lateral septum peaks immediately before feeding (during food anticipatory activity) in rabbit pups (Olivo et al., 2017) and gamma-rhythmic input to the LHA from the LS evokes food approach behavior (Carus-Cadavieco et al., 2017). In addition, the LS can bidirectionally modulate food intake via several feeding-relevant neuropeptides, including glucagon-like peptide-1, ghrelin, and urocortin (Maske et al., 2018; Terrill et al., 2016; Wang and Kotz, 2002). These feeding-associated results are consistent with the contribution of the CA1v-to-LS pathway in promoting spatial memory in the context of foraging but not in the escape-based version of the visuospatial task. Given that the rats were food or water-restricted for the respective food- and water-seeking memory tasks, it is likely that a CA1v-to-LS pathway potentiates visuospatial memory in response to changes in internal state. Moreover, this internal state modulation of CA1v-to-LS control of appetitive behavior is specific to foraging-related memory as disconnection of the circuit did not influence motivated operant responding for the same reinforcer (sucrose) used in the food-seeking spatial memory task.

Recent findings demonstrate that DREADDs-mediated disconnection of a HPCv to LS pathway (from CA3v and DGv) increases food intake in mice, with opposite/hyperphagic effects observed following activation of this pathway (Sweeney and Yang, 2015). Here we observe no differences in food intake nor meal parameters following DREADDs-mediated acute silencing of the CA1v to LS in rats with unlimited access to food and water, therefore establishing a role for this pathway in foraging-relevant memory processes under conditions that are more relevant to normal mammalian behavior, where food and water are not freely available. HPCv projections to the LS have also been shown to influence anxiety-like behaviors (Parfitt et al., 2017; Trent and Menard, 2010). However, neither our acute nor chronic disconnection model of the CA1v-to-LS pathway affected these behaviors. Given that these studies targeted the HPCv either independently of subregion or targeted CA3v and DGv, it is probable that functions of HPCv-to-LS pathways differ according to subregional specificity. In fact, a recent study demonstrated opposite effects of inhibition of CA1v and CA3v projections to the LS on approach-avoidance conflict resolution (Yeates et al., 2022).

This work focused on the contribution of an HPCv-LS neural pathway to foraging-related memory, yet it is likely that the HPCd also partakes in appetitive spatial memory. In fact, lesions of the HPCd were reported to impair performance in a reinforced radial arm maze procedure(Gaskin et al., 2009) and simultaneous recordings from HPCd and LS cells revealed comparable activity patterns in these two regions during spatial navigation towards a food reward (Wirtshafter and Wilson, 2020). Foraging-relevant memory processing in the HPCd may also engage the LS, as oscillatory patterns from HPCd neurons are predictive of interstitial peripheral glucose dynamics via projections to the LS (Tingley et al., 2021). In addition, the feeding-related neuropeptide MCH was shown to promote spatial learning and memory consolidation by strengthening excitatory signaling within a HPCd-LS neural pathway (Liu et al., 2022). The extent to which HPCv and HPCd inputs to the LS independently promote foraging-related memory remains to be elucidated.

The use of two different but complementary approaches for neural pathway-specific disconnection allowed us to test at the levels of learning and memory separately. For example, the use of a reversible (DREADDs) disconnection procedure allowed us to evaluate the effect of CA1v to LS disconnection during the memory probe only, with circuit intact during training (CNO was not administered at any point during training). In contrast, the chronic (caspase) disconnection procedure was performed before training and therefore could have affected learning if CA1v to LS pathways were required for the learning of the task. However, our results showed that CA1v to LS disconnection did not impair learning, but rather, only memory retention performance in the probe test conducted days after the end of training. Thus, we can now confirm based on two complementary levels of analysis that this pathway is involved in longer-term memory recall but not learning/acquisition, which is consistent with recent evidence demonstrating that complete HPCv lesions impair retrieval (probe performance), but not acquisition (learning) in a Morris water maze (Hauser et al., 2020). A limitation of the present study is that we did not evaluate ‘gain of function’ of CA1v->LS signaling on appetitive spatial memory. However, we note that such gain of function approaches are more commonly utilized with models of memory dysfunction. Thus, an important area for follow-up research is to investigate whether enhanced CA1v->LS signaling (e.g., via excitatory pathway-specific DREADDs) can augment memory function in models where foraging-related memory is impaired.

While the use of aversive reinforcement-based spatial memory tasks is predominant in the field, the radial arm maze has also been used to investigate food-motivated spatial memory (Olton, 1987). An advantage of the present food-reinforced Barnes maze apparatus-based approach over the radial arm maze procedure is that we have developed parallel procedures that use the same apparatus and visuospatial cues to examine water- and escape-motivated spatial memory, thus offering a unique arsenal to assess the selectivity of the circuit based on multiple reinforcement modalities (food vs. water vs. escape/aversive). Future research is needed to determine whether CA1v to LS signaling is also involved in spatial working memory based on appetitive and/or aversive reinforcement.

Utilizing dual viral approaches and functional immediate early gene mapping analyses, our findings confirm that the CA1v-LS pathway projects downstream to the LHA. This region was the most robust forebrain downstream target of CA1v->LS signaling, and this multisynaptic pathway was corroborated by an additional monosynaptic tracing approach that combined retrograde tracing (FG in LHA) and anterograde tracing (AAV1-GFP in CA1v) to confirm a connection relay in the LS. The identification of the majority of LS neurons receiving input from the CA1v as GABAergic, along with the resulting increase in medial LHA c-Fos expression following acute disconnection of the CA1v-to-LS pathway suggests that this chemogenetic intervention prevents medial LHA local inhibition of excitatory neurons. This is consistent with previous findings reporting disinhibition of LHA GABAergic neurons by projecting GABAergic neurons in the context of feeding (O’Connor et al., 2015; Sweeney and Yang, 2015).

We have previously found that the CA1v sends direct projections to the LHA to modulate food intake and meal size control by the orexigenic gastric hormone, ghrelin (Hsu et al., 2015; Suarez et al., 2020). In the present study, we highlight a novel multisynaptic pathway connecting the CA1v to the medial LHA through the LS to promote foraging-related spatial memory. These findings suggest that the CA1v is functionally-connected to the LHA through direct and indirect projections to influence food intake and appetitive spatial memory, respectively. Interestingly, the direct CA1v-LHA pathway was shown to target orexin neurons whereas the current study finds that although medial LHA orexin neurons are engaged during the appetitive spatial memory probe, this activation is not influenced by acute disconnection of the CA1v-LS-LHA indirect pathway. Thus, these parallel pathways may be promoting differential feeding-relevant behaviors through different neuronal populations in the LHA, with the indirect pathway mediating appetitive memory and the direct pathway mediating food consumption.

Collective results identify a CA1v to LS pathway involved in food- and water-motivated spatial memory, but not escape-motivated spatial memory. Furthermore, the selectivity of this pathway to appetitive spatial memory is supported by data showing that neither chronic nor reversible disruption of CA1v to LS signaling influenced various other behavioral outcomes, including food intake, anxiety-like behavior, locomotor activity, and nonspatial HPC-dependent appetitive memory. We also systematically characterized collateral (mPFC) and second-order (LHA) projections of this pathway that are not functional, and functional (respectively), for appetitive spatial memory. Present results shed light on the neural systems underlying complex learned and motivated behaviors that require functional connections between cognitive and feeding-relevant substrates.

## ACKNOWLEDGEMENTS

This work was supported by a Fonds de Recherche du Québec Postdoctoral Fellowship to LDS (315201) and by National Institute of Diabetes and Digestive and Kidney Diseases grants: DK104897 to SEK, DK118944 to CML, DK116558 to ANS, and DK118000 to EEN. Clozapine-N-Oxide was kindly provided by the National Institute of Mental Health. The authors are grateful to the Kanoski lab undergraduate research assistants for their assistance in behavioral experiments and histology.

## AUTHOR CONTRIBUTIONS

LDS, CML, EAD and SEK conceived and performed experiments and wrote the manuscript. LTL, KS, AGB, MEK, IHG, ANS, KND and AMC performed experiments. EEN provided expertise. JDH performed experiments and provided expertise.

## RESOURCE AVAILABILITY

### Lead contact

Further information and requests for resources and reagents should be directed to and will be fulfilled by the lead contact, Scott Kanoski.

### Materials availability

This study did not generate new unique reagents

### Data and code availability

- Original data have been deposited at the Open Science Framework Repository and are publicly available as of the date of publication. Accession number is listed in the key resources table.
- Original code has been deposited to the Open Science Framework Repository and are publicly available as of the date of publication. Accession number is listed in the key resources table.
- Any additional information required to reanalyse the data reported in this paper is available from the lead contact upon request.

## EXPERIMENTAL MODEL AND SUBJECT DETAILS

Adult male Sprague-Dawley rats (Envigo, 250-275g on arrival) were individually housed in hanging wire cages with *ad libitum* access (except where noted) to water and chow (LabDiet 5001, LabDiet, St. Louis, MO) on a 12h:12h reverse light/dark cycle. All procedures were approved by the University of Southern California Institute of Animal Care and Use Committee. Sample size for all behavioral and neuroanatomical experiments was based on the authors’ extensive experience with the procedures and statistical power analyses (described below).

## METHOD DETAILS

### Appetitive food seeking spatial memory task

To test visuospatial learning and memory for food reinforcement, we developed a spatial food seeking task modified from the traditional Barnes maze procedure (Barnes, 1979). Throughout this paradigm, animals were maintained at 85% free-feeding body weight. In contrast to the traditional Barnes maze where an animal uses the spatial cues to escape mildly aversive stimuli in the environment (e.g. bright light and loud sound), this task utilizes food as motivation, such that each hidden tunnel contained five 45mg sucrose pellets (cat. no. F0023, Bio-Serv, Flemington, NJ).

The procedure involves an elevated gray circular platform (diameter: 122cm, height: 140cm) consisting of 18 uniform holes (9.5cm diameter) spaced every twenty degrees around the outside edge. Under one of the holes is a hidden tunnel (38.73cm L x 11.43cm W x 7.62cm D and a 5.08cm step). Surrounding the table are distinct spatial cues on the wall (e.g., holiday lights, colorful shapes, stuffed unicorn) that are readily visible to the animal. Additionally, a quiet white noise (60dB) was used to drown out background noise and floor lamps were used for low-level ambient lighting.

On the first day, each animal underwent a habituation session consisting of 1min inside a transport bin under the table, 2min underneath the start box in the center of the table, and 3 min inside the hidden tunnel with five sucrose pellets. During training, each rat was assigned a specific escape hole according to the position of the spatial cues with the location counterbalanced across groups. Before each trial, animals were placed in the start box for 30 seconds. Animals were trained over the course of two 5-min trials per day for four days (five days in contralesional experiments) to learn to use the spatial cues in order to locate the correct hole with the hidden tunnel with sucrose pellets. After finding the correct hole and entering the hidden tunnel during each 5min training session, animals were allowed to remain in the hidden tunnel for 1 minute, and consistently consumed all five sucrose pellets at this time. Learning during training was scored via animal head location tracking by AnyMaze Behavior Tracking Software (Stoelting Co., Wood Dale, IL). The incorrect hole investigations prior to finding the correct hole with sucrose pellets (“errors before correct hole”) as well as time to finding the correct hole (“latency to correct hole”) were calculated as an average of the two trials per day and examined across days of training. After the conclusion of training, rats had a two-day break where no procedures occurred, then were tested in a single 2min memory probe in which the hidden tunnel and sucrose pellets were removed.

In the event that a rat did not find the correct hole during the 3min training session, the experimenter gently guided the rat to the correct hole and were allowed to crawl in and then complete the training in the same fashion as rats that found the hole on their own. During training sucrose pellets were taped underneath all of the holes to preclude olfactory-based search strategies during training. Importantly, the memory probe test does not include any sucrose pellets in the apparatus at all, and therefore there is no possibility of sucrose odor cues to influence memory probe performance.

In the memory probe, the ratio of the correct hole plus adjacent hole investigations over the total number of hole investigations were calculated via animal head location tracking by AnyMaze Behavior Tracking Software. This dependent variable was selected a priori instead of correct hole only investigations based on the spatial resolution of rodent hippocampal place cells which ranges from inches to meters depending on the subregion targeted (Kjelstrup et al., 2008) and recent findings demonstrating that taste-responsive hippocampal neurons exhibit weaker spatial selectivity in comparison to non-taste-responsive hippocampal neurons (Herzog et al., 2019). This biological range is better represented by a geometric plane that includes 3/18 circumference space versus 1/18 as we have previously reported (Suarez et al., 2018b).

When establishing and refining the food-reinforced spatial memory procedure in multiple cohorts of rats, our preliminary data revealed that memory probe effects in normal/control animals are either stronger in the 1^st^ minute, or in the 1^st^ two minutes combined (but not the 2^nd^ minute alone), with some cohorts showing larger paradigm effects in minute 1, and others in minutes 1+2. Thus, for all experiments presented we analyzed both the 1^st^ minute and the 1^st^ two minutes combined based on time at which paradigm effect is the strongest.

### Intracranial injections

Rats were anesthetized via an intramuscular injection of an anesthesia cocktail (ketamine 90mg/kg body weight [BW], xylazine, 2.8mg/kg BW and acepromazine and 0.72mg/kg BW) followed by a pre-operative, subcutaneous injection of analgesic (ketoprofen, 5mg/kg BW). Post-operative analgesic (subcutaneous injection of ketoprofen, 5mg/kg BW) was administered once per day for 3 days following surgery. The surgical site was shaved and prepped with iodine and ethanol swabs, and animals were placed in a stereotaxic apparatus for stereotaxic injections. NMDA or viruses were delivered using a microinfusion pump (Harvard Apparatus, Cambridge, MA) connected to a 33-gauge microsyringe injector attached to a PE20 catheter and Hamilton syringe. Flow rate was calibrated and set to 83.3nl/sec. Injectors were left in place for 2min post-injection to allow for complete delivery of the infusate. Specific viruses/drugs, coordinates, and injection volumes for procedures are detailed below. Following the completion of all injections, incision sites were closed using either surgical staples, or in the case of subsequent placement of an indwelling cannula, simple interrupted sutures. Upon recovery from anesthesia and return to dorsal recumbency, animals were returned to the home cage. All behavioral procedures occurred 21 days after injections to allow for complete transduction and expression of the viruses, or complete lesioning drugs. General intracranial injection procedures were followed for all injection procedures below.

### Fiber photometry

To record bulk calcium-dependent activity in the HPCv, one group of animals (n=6) received a bilateral injection of a virus driving expression of the calcium indicator GCaMP7s (AAV9-hSyn-GCaMP7s-WPRE; Addgene; 300nL per side) targeting the CA1v at the following stereotaxic coordinates: −4.9mm AP, +/−4.8mm ML, defined at bregma and −7.8mm DV, defined at skull site. A fiber-optic cannula (Doric Lenses Inc, Quebec, Canada) was implanted at the injection site and affixed to the skull with jeweler’s screws, instant adhesive glue, and dental cement.

Photometry recording sessions occurred during the probe trial of the appetitive spatial food seeking memory task (when no food is present) and lasted 3 minutes (1 min acclimatation and 2 min probe trial). Photometry signal was acquired using the Neurophotometrics fiber photometry system (Neurophotometrics, San Diego, CA) at a sampling frequency of 40Hz and calcium-dependent signal (470nm) was corrected in MatLab by subtracting the isosbestic (415nm) value and fitting the result to a biexponential curve. Corrected fluorescence signal was then normalized within each rat by calculating the ΔF/F using the average fluorescence signal for the entire recording. The data was paired with real-time localization of the animal obtained from the AnyMaze Behavior Tracking Software (Stoelting, Wood Dale, IL) using an original MatLab code and each investigation at least 5 seconds apart were isolated for analysis. Z-scores were obtained by normalizing ΔF/F up to 5 seconds following an investigation to the ΔF/F at the onset of the investigation (time 0s).

### Chronic lesions of the HPCv

Lesion animals (n=18) received bilateral excitotoxic HPCv lesions via intracranial injections of N-methyl-d-aspartate (NMDA, Sigma, St-Louis, MO; 4ug in 200nL; 100uL per hemisphere) at the following coordinates at three different sites along the rostro-caudal extent of the HPCv: [1] −4.8mm AP, +/−5.0mm ML, −7.5mm DV, [2] −5.5mm AP, +/−4.5mm ML, −7.0mm DV, and [3] −6.1mm AP, +/−4.8mm ML, −7.0mm DV with control animals (n=11) receiving vehicle saline in the same location. The reference points for AP and ML coordinates were defined at bregma, and the reference point for the DV coordinate was defined at the skull surface at the target site.

Bilateral HPCv lesion brains were histologically evaluated for the correct placement of lesions in 1 out of 5 series of brain tissue sections. Neurons were visualized using immunohistochemistry for the neuron specific antigen NeuN (see Immunohistochemistry). The extent of NMDA lesions was determined postmortem by immunohistochemical detection of the neuronal marker NeuN. Rats showing pronounced (∼65%) loss of NeuN labeling within the target region compared with the representative NeuN expression following sham injections in the control group (average HPCv count of NeuN+ cells in control group) were included in data analysis.

### Acute and chronic disconnection models

Cre-dependent dual viral strategies were used to generate the following groups: [1] acute chemogenetic disconnection of the CA1v to LS neural pathway (DREADDs), [2] chronic disconnection of the CA1v to LS neural pathway (caspase lesion-induced), and [3] a common/shared control group for these two disconnection strategies. Regardless of group, all animals received a bilateral AAV retro-cre injection in the LS (AAV2[retro]-hSYN1-EGFP-2A-iCre-WPRE; Vector BioLabs; 200nL per side) at the following stereotaxic coordinates: +0.84mm AP, +/−0.5mm ML, −4.8mm DV, all defined at bregma. According to experimental group, animals received a different virus delivered to the CA1v subregion of the HPCv at the following coordinates: −4.9mm AP, +/-4.8mm ML, defined at bregma and −7.8mm DV, defined at skull site.

[1] *DREADDs group for reversible inactivation of CA1v to LS:* To allow reversible chemogenetic inactivation of the CA1v to LS neural pathway, one group of animals received a bilateral CA1v injection of a cre-dependent virus to drive expression of inhibitory designer receptors activated by designer drugs (DREADDs), (AAV-Flex-hM4Di-mCherry; Addgene; 200nL per side). This dual viral strategy drives expression of inhibitory DREADDs exclusively in LS-projecting CA1v neurons, which enables acute inactivation of these neurons by injection of the DREADDs ligand, clozapine-N-oxide (CNO), at the time of behavioral testing.
[2] *Caspase group for chronic inactivation of CA1v to LS:* To allow chronic disconnection of the CA1v to LS neural pathway, a second group of animals received a bilateral CA1v cre-dependent caspase virus (AAV1-Flex-taCasp3-TEVp; UNC Vector Core; 200nL per side) mixed with a cre-dependent reporter virus (AAV-CAG-Flex-tdTomato-WPRE-bGH, UPenn Vector Core; 200nL per side) for histological verification. This dual viral strategy drives expression of the apoptosis-mediator molecule caspase exclusively in LS-projecting CA1v neurons, which induces apoptotic cell death in these neurons while leaving other CA1v neurons intact.
[3] *Common control:* A common control group was used for the DREADDs and caspase groups. This allowed us to reduce the number of animals needed for the same experimental objective. Thus, all animals received an indwelling cannula and ICV injections of CNO as described below.

Immediately following viral injections, all animals were surgically implanted with a unilateral indwelling intracerebroventricular (ICV) cannula (26-gauge, Plastics One, Roanoke, VA) targeting the lateral ventricle (VL). Cannulae were implanted and affixed to the skull with jeweler’s screws and dental cement at the following stereotaxic coordinates: −0.9mm AP defined at bregma, +1.8mm ML defined at bregma, −2.6mm DV defined at skull surface at site.

### Validation of inhibitory DREADDs expression

Cre-dependent DREADD expression targeting LS-projecting neurons in CA1v was evaluated based on localization of the fluorescent reporter mCherry. Immunohistochemistry for red fluorescent protein (RFP) was conducted to amplify the mCherry signal (see Immunohistochemistry). Only animals with mCherry expression restricted within CA1v neurons were included in subsequent behavioral analyses.The same approach was used to validation of inhibitory DREADDs expression for the CA1v-mPFC experiment.

### Validation of caspase-mediated disconnection

The general approach for neuronal apoptosis due to activation of cre-dependent caspase targeting LS-projecting neurons in CA1v was validated in a separate cohort (n=8) based on reduction of a cre-dependent fluorescent reporter tdTomato due to neuron cell death compared with controls. The caspase group brains (which received a retro-cre virus in the LS in conjunction with an injection of cre-dependent caspase virus mixed with a cre-dependent virus that drives a tdTomato fluorescent reporter in the CA1v) were compared to control brains (that received a retro-cre virus in the LS in conjunction with only a cre-dependent that drives a tdTomato fluorescent reporter in the CA1v, which was equivalently diluted to match the caspase injections). In both groups, immunohistochemistry for red fluorescent protein (RFP) was conducted to amplify the tdTomato signal (see Immunohistochemistry). Confirmation of successful caspase-driven lesions in the CA1v using this approach was evaluated by reduced tdTomato fluorescence in comparison to control animals, with the expression of CA1v tdTomato expression in the caspase group less than 15% of that observed in controls, on average. While this cohort allowed for successful validation of the pathway-specific disconnection caspase approach, to improve the viability of the animals for long-term behavioral analyses, the 3^rd^ AAV with the fluorescent reporter (cre-dependent tdTomato AAV) was omitted for cohorts undergoing behavioral analyses. For these groups, histological confirmation for inclusion in subsequent statistical analyses was based on identification of injections sites confined with the LS and CA1v, as observed using darkfield microscopy.

### Validation of ICV cannula placement

Placement for the VL cannula was verified by elevation of blood glucose resulting from an injection of 210μg (2μL) of 5-thio-D-glucose (5TG) using an injector that extended 2mm beyond the end of the guide cannula (Ritter et al., 1981). A post-injection elevation of at least 100% of baseline glycemia was required for subject inclusion. Animals that did not pass the 5TG test were retested with an injector that extended 2.5mm beyond the end of the guide cannula and, upon passing the 5TG test, were subsequently injected using a 2.5mm injector instead of a 2mm injector for the remainder of the study.

### ICV CNO infusion

Administration route for CNO (RTI International) and dosage were determined based on previous publications demonstrating reliable DREADDs activation (Hsu et al., 2018a; Noble et al., 2018; Terrill et al., 2020). Prior to behavioral testing where noted, all animals received an ICV 18mmol infusion (2uL total volume) of the DREADDs ligand clozapine N-oxide (CNO), rendering only DREADDs animals chemogenetically inactivated. CNO injections were hand delivered using a 33-gauge microsyringe injector attached to a PE20 catheter and Hamilton syringe through the indwelling guide cannula. Injectors were left in place for 30sec to allow for complete delivery of the CNO.

### Appetitive water seeking spatial memory task

We modified our spatial food seeking task to utilize water reward instead of food reward. Throughout this paradigm, animals were water restricted and received 90-min access to water daily (given at least 1hr after the termination of behavioral procedures each day). In addition to the habituation parameters described above, animals in this task were subjected to two 10-min habituation sessions in the hidden tunnel with a full water dish prior to training, which allowed them to become accustomed to reliably drinking from the water dish in the hidden tunnel. Training procedures were the same as for the food-based task above, except instead of sucrose pellets in the hidden tunnel, a water dish containing ∼100 mL of water was placed in the back of the hidden tunnel during training, and animals were allowed to remain in the hidden tunnel for 2min (instead of 1min, as above). Each animal consumed a minimum of 2mL of water during this time for the training sessions. This adapted procedure allowed us to test spatial learning and memory motivated by water reward using similar apparatus and stimulus conditions to the food-reinforced task.

### Aversive spatial memory escape task (Barnes maze)

To test visuospatial learning and memory for escape-based reinforcement, we used a modified traditional Barnes maze procedure, which is a visuospatial escape task (Barnes, 1979). Procedures were the same as above (appetitive spatial memory food seeking and water seeking tasks, using the same apparatus, in the same room, and with the same visuospatial cues) aside from the omission of the sucrose pellets or water dish in the hidden tunnel, the presence of mildly aversive bright (120W) overhead lighting instead of dim ambient lighting, and a mildly aversive loud white noise (75dB) instead of a quiet white noise (60dB). This allowed us to test spatial learning and memory motivated by escape from aversive stimuli in a nearly identical procedure to our spatial foraging test for learning and memory motivated by palatable food or water consumption.

### Identification of CA1v to LS collateral targets

Collateral targets of the CA1v to LS neural pathway were identified using a dual viral approach where a retrograde vector was injected into the LS (AAV2retro-hSyn1-eGFP-2A-iCre-WPRE; Vector BioLabs; 200nL per side), and a Cre-dependent anterograde vector (AAV1-CAG-Flex-tdTomato-WPRE-bGH, UPenn Vector Core; 200nL per side) was injected in the CA1v. This latter injection drives tdTomato transgene expression in CA1v neurons that project to the LS, which allows for brain-wide analyses of collateral targets. After 3 weeks of incubation time to allow for complete transduction and expression of the viruses, brains were collected, immunohistochemically processed, and imaged as described below.

### Acute CA1v to mPFC disconnection

To functionally disconnect the CA1v to mPFC pathway (diagram of approach in Fig. 3C) a group of animals all received bilateral AAV retro-cre injection in the mPFC (AAV2[retro]-hSYN1-EGFP-2A-iCre-WPRE; Vector BioLabs; 200nL per side) at the following stereotaxic coordinates: +2.7mm AP, +/− 0.5mm ML and −4.3mm DV, all defined at bregma. The same animals also received bilateral infusion of a cre-dependent inhibitory DREADDs (AAV-Flex-hM4Di-mCherry; Addgene; 200nL per side) in the CA1v at the following stereotaxic coordinates: −4.9mm AP, +/−4.8mm ML, defined at bregma and −7.8mm DV, defined at skull site. Finally, all these animals were additionally implanted with a unilateral indwelling ICV cannula as described for the previous cohorts at the following stereotaxic coordinates: −0.9mm AP defined at bregma, +1.8mm ML defined at bregma, −2.6mm DV defined at skull surface at site. Prior to the memory probe (1h), animals received an ICV 18mmol infusion of CNO (n=8) or its vehicle (aCSF; n=6).

### Food intake and body weight

For the day prior to surgery (day 0) and for two weeks thereafter, 24h chow intake was measured daily just prior to dark cycle onset to determine effects of experimental procedures on food intake. Spillage was accounted for daily by collecting crumbs onto Techboard paper placed under the cages of each animal. Additionally, animals were weighed daily just prior to dark cycle onset to determine effects of experimental procedures on body weight.

### Meal pattern analysis

Meal size, meal frequency, meal duration, and cumulative 24h food intake were measured using Biodaq automated food intake monitors (Research Diets, New Brunswick, NJ). Rats were acclimated to the Biodaq on ad libitum chow for 3 days. Caspase group (n=7) vs. control group (n=7) feeding behavior data were collected over a 5-day period (untreated, between-subjects design). The DREADDs group was tested using a 2-treatment within-subjects design. Rats (n=8) received 18mmol infusion of ICV CNO (2uL total volume) or vehicle (aCSF, 33% dimethyl sulfoxide in artificial cerebrospinal fluid) 1h prior to lights off and 24h feeding behavior was measured. Meal parameters were set at minimum meal size=0.2g and maximum intermeal interval=600s.

### Operant responding for sucrose rewards

Rats expressing inhibitory DREADDs in CA1v-to-LS projections and implanted with an ICV cannula (n=6) were trained to lever press for 45mg sucrose pellets (cat. no. F0023, Bio-Serv) over the course of 6 days with a 1h session each day (2 days of fixed ratio 1 with autoshaping procedure, 2 days of fixed ratio 1 and 2 days of fixed ratio 3 reinforcement schedule) in operant conditioning boxes (Med Associates Inc, St. Albans, VT). Over two treatment days separated by 48h, rats were placed back in the operant chambers to lever press for sucrose under a progressive ratio reinforcement schedule, 1h following ICV infusion (2uL) of 18mmol CNO or aCSF, with both treatments administered to each rat in a counter-balanced manner (within-subject). The response requirement increased progressively using the following formula: F(i) = 5ê0.2i-5, where F(i) is the number of lever presses required for the next pellet at i = pellet number and the breakpoint was defined as the final completed lever press requirement that preceded a 20-min period without earning a reinforcer, as described previously (Hsu et al., 2015b).

### Social transmission of food preference (STFP)

To examine food-related memory based on social- and olfactory-based cues, we utilized the social transmission of food preference (STFP) task and adapted protocols from (Countryman and Gold, 2007; Countryman et al., 2005; Hsu et al., 2018b). Briefly, untreated normal adult rats were designated as ‘Demonstrators’, while experimental groups were designated as ‘Observers’. Demonstrators and Observers were habituated to a powdered rodent chow [LabDiet 5001 (ground pellets), LabDiet, St. Louis, MO] overnight. 24h later, Observers were individually paired with demonstrators and habituated to social interaction, where rat pairs were placed in a social interaction arena (23.5cm W × 44.45cm L × 27cm H clear plastic bin with Sani-chip bedding) and allowed to interact for 30min. Both Observers and Demonstrators were returned to their home cages and food was withheld for 23hr prior to the social interaction. For the social interaction, Demonstrators were given the opportunity to consume one of two flavors of powdered chow (flavored with 2% marjoram or 0.5% thyme; counterbalanced according to group assignments) for 30min in a room separate from Observers. Our pilot studies and previous published work (Galef and Whiskin, 2003; Galef et al., 2005) showed that rats equally prefer these flavors of chow. The Demonstrator rat was then placed in the social interaction arena with the Observer rat, and the pairs were allowed to socially interact for 30min. Observers were then returned to their home cage and allowed to eat *ad libitum* for 1h and then food was removed. The following day, the 23h food-deprived Observer animals were given a home cage food preference test for either the flavor of chow paired with the Demonstrator animal, or a novel, unpaired flavor of chow that is a flavor that was not given to the Demonstrator animal (2% marjoram vs. 0.5% thyme; counterbalanced according to group assignments). All animals received an ICV 18mmol infusion of CNO (2uL total volume) 1h prior to the social interaction session, rendering only DREADDs animals chemogenetically inactivated by CNO. Food intake (1h) was recorded with spillage accounted for by collecting crumbs onto Techboard paper that was placed under the cages of each animal. The % preference for the paired flavor was calculated as: 100*Demonstrator-paired flavored chow intake/Demonstrator + Novel flavored chow intake. In this procedure, normal untreated animals learn to prefer the Demonstrator paired flavor based on social interaction and smelling the breath of the Demonstrator rat (Countryman and Gold, 2007; Countryman et al., 2005; Galef and Whiskin, 2003; Galef et al., 2005; Hsu et al., 2018b).

### Zero maze

The zero maze behavioral paradigm was used to evaluate anxiety-like behavior. The zero maze apparatus used was an elevated circular track, divided into four equal length sections. Two zones were open with 3 cm high curbs (‘open zones’), whereas the two other zones were closed with 17.5 cm high walls (‘closed zones’). All animals received an ICV 18mmol infusion of CNO (2uL total volume) 1h prior to testing, rendering only DREADDs animals chemogenetically inactivated by CNO. Behavioral testing was performed during the light cycle. Animals were placed in the maze for a single, 5min trial in which the location of the center of the animal’s body was measured by AnyMaze Behavior Tracking Software (Stoelting). The apparatus was cleaned with 10% ethanol in between animals. During the trial, the number of open zone entries and total time spent in open sections (defined as body center in open sections) were measured, which are each indicators of anxiety-like behavior in this procedure.

### Open field

An open field test was used to evaluate general levels of locomotor activity. The apparatus used for the open field test was an opaque gray plastic bin (60cm × 56cm), which was positioned on a flat table in an isolated room with a camera directly above the center of the apparatus. Desk lamps were positioned to provide indirect lighting to all corners of the maze such that the lighting in the box uniformly measured 30 lux throughout. All animals received an ICV 18mmol infusion of CNO (2uL total volume) 1h prior to testing rendering only DREADDs animals chemogenetically inactivated by CNO. Behavioral testing began at dark onset. At the start of the 10min test, each animal was placed in the open field apparatus in the same corner facing the center of the maze. The location of the center of the animal’s body was measured with the AnyMaze Behavior Tracking Software (Stoelting). Total distance traveled was measured by tracking movement from the center of the animal’s body throughout the test.

### c-Fos + FISH neurotransmitter phenotyping

Activation of LS neurons from the spatial memory probe test was confirmed with c-Fos immunoreactivity analyses. Animals receiving the DREADDs-mediated treatment for reversible inactivation of CA1v to LS signaling underwent training in the appetitive food seeking memory task (conducted as described above). One hour before the memory probe, animals received ICV injection of CNO (n=4) or vehicle (aCSF; n=4). Animals were tested in the memory probe and perfused with tissue harvest 90 min later. Fluorescent *in situ* hybridization analyses for c-Fos (cat. no. 403591, ACD, Newark, CA), ChAT (cat. no. 430111, ACD), GAD1 (cat. no. 316401, ACD), and vGLUT2 (cat. no. 317011, ACD) were performed, and co-localization of c-Fos positive cells and individual neurotransmitter markers was quantitatively analyzed for a subset of rats (n=3) in the vehicle treated group.

### Identification of second-order targets

To identify second order targets of CA1v to LS-projecting neurons, animals received a bilateral injection of a transsynaptic Cre-inducing anterograde vector into CA1v (AAV1-hSyn-Cre-WPRE-hGH, UPenn Vectore Core; 200nL per side; coordinates as above) that drives expression of Cre in both first-order (CA1v injection site) and 2^nd^-order (but not 3^rd^-order) neurons via transsynaptic virion release (Suarez et al., 2018b; Zingg et al., 2017). This was combined with a bilateral injection of a Cre-dependent anterograde vector in the LS (AAV1-CAG-Flex-tdTomato-WPRE-bGH, UPenn Vector Core; 200nL per side, coordinates as above). This latter injection allows for anterograde tracing from 1^st^-order LS targets receiving synaptic input from CA1v. After 3 weeks of incubation time to allow for complete transduction and expression of the viruses, brains were collected, immunohistochemically processed, and imaged as described below.

Data were entered using a custom built data-entry platform (Axiome C, created by JDH) built around Microsoft Excel software and designed to facilitate entry of data points for all gray matter regions across their atlas levels as described in a rat brain reference atlas: Brain Maps 4.0 (Swanson, 2018). The Axiome C approach was used previously to facilitate the analysis of brain gene-expression data (Hahn et al., 2019). An ordinal scale, ranging from 0 (absent) to 7 (very strong), was used to record the qualitative weight of anterograde labeling. An average value was then obtained for each region across its atlas levels for which data were available. These data are summarized graphically for a representative animal on a brain flatmap summary diagram (adapted from (Hahn et al., 2021)).

The LHA was identified as a primary downstream target of the CA1v to LS pathway from the analyses described above. In order to further confirm this conclusion, an additional co-injection monosynaptic neural tracing strategy (COIN; (Suarez et al., 2018b; Thompson and Swanson, 2010)) was conducted. Animals (n=6) received unilateral pressure injections of AAV1-GFP (AAV1-hSyn-eGFP-WPRE-bGH; Vector BioLabs; anterograde tracer) targeting the CA1v and, in the same animal, fluorogold (Fluorochrome; retrograde tracer) targeting the LHA (injection coordinates described above). Following a 10-day incubation period, animals were fixation-perfused and tissue was collected and processed as described below. Injection sites were confined to the CA1v (AAV1) and LHA (FG) in the same animal (“double hits”, n=4). Appositions between FG and GFP were examined in the LS in these animals by visualizing native GFP and FG fluorescence and the percentage of FG cells in direct contact with GFP CA1v-originating axons was quantified (n=4).

To verify if acute silencing of the CA1v to LS neuronal pathway reduces neuronal activity in the LHA, staining against c-Fos was performed on coronal LHA slices from animals with expression of inhibitory DREADDs in CA1v to LS neurons and infused with CNO (n=4) or vehicle (aCSF; n=4) 1h prior to the memory probe. Counts of c-Fos+ cells were done in the 2 subregions of the LHA found to receive the highest density of projections from CA1v neurons (LHAjd and LHAjv). In another series of LHA coronal sections from the same animals, counts of c-Fos+ cells were done in orexin+ or MCH+ neurons (see Immunohistochemistry).

Counts of c-Fos+ cells were also done in LHA coronal sections from rats treated with CNO (n=6) and aCSF (n=4) lacking expression of inhibitory DREADDs as a control experiment.

### Immunohistochemistry

Rats were anesthetized via an intramuscular injection of an anesthesia cocktail (ketamine 90mg/kg BW xylazine, 2.8mg/kg BW and acepromazine and 0.72mg/kg BW) then transcardially perfused with 0.9% sterile saline (pH 7.4) followed by 4% paraformaldehyde (PFA) in 0.1M borate buffer (pH 9.5; PFA). Brains were dissected from the skull and post-fixed in PFA with 15% sucrose for 24h, then flash frozen in isopentane cooled in dry ice. Brains were sectioned to 30µm thickness on a freezing microtome. Sections were collected in 5-series and stored in antifreeze solution at −20°C until further processing. General fluorescence IHC labeling procedures were performed as follows. The antibodies and dilutions that were used are as follows: [1] For lesion histology using the neuron-specific protein NeuN, the rabbit anti-NeuN primary antibody (1:1000, Abcam, Boston, MA) was used followed by a donkey anti-rabbit conjugated to AF488 (1:500, Jackson Immunoresearch, West Grove, PA). [2] To amplify the native tdTomato signal for neuroanatomical tracing or DREADDs histology, the rabbit anti-RFP primary antibody (1:2000, RRID: AB_2209751, Rockland, Limerick, PA) was used followed by a donkey anti-rabbit conjugated to Cy3 (1:500, Jackson Immunoresearch). [3] To amplify the native GFP signal for LS injection site histology, the chicken anti-GFP primary antibody (1:500, Abcam) was used followed by a donkey anti-chicken secondary antibody conjugated to AF488 (1:500, Jackson Immunoresearch). [4] To validate that the acute disconnection approach reduced neuronal activity in the LS and LHA, the mouse anti-c-Fos primary antibody (1:1000, Ab208942, Abcam) was used followed by a donkey anti-mouse secondary antibody conjugated to AF647 (1:500, Jackson Immunoresearch). To identify the neuropeptide population in which LHA c-Fos is expressed following the foraging spatial memory probe, the rabbit anti-orexin (1:5000, AB3704, EMD Millipore, Burlington, MA) or anti-MCH (1:10000, H_070_47, Phoenix Pharmaceuticals, Burlingame, CA) primary antibody was also used followed by a donkey anti-rabbit secondary antibody conjugated to AF488 (1:500, Jackson Immunoresearch).

Antibodies were prepared in 0.02M potassium phosphate buffered saline (KPBS) solution containing 0.2% bovine serum albumin and 0.3% Triton X-100 at 4°C overnight. After thorough washing with 0.02M KPBS, sections were incubated at 4°C overnight in secondary antibody solution. Sections were mounted and coverslipped using 50% glycerol in 0.02 M KPBS and the edges were sealed with clear nail polish. Photomicrographs were acquired using a Nikon 80i (Nikon DSQI1,1280X1024 resolution, 1.45 megapixel) under epifluorescence or darkfield illumination.

Lesions and virus expression were quantified in one out of five series of brain tissue sections and analyses were performed in sections from Swanson Brain Atlas level 34-36 (Swanson, 2018). The inclusion / exclusion criteria are described above in the methods of each of these experiments.

## QUANTIFICATION AND STATISTICAL ANALYSIS

Data are expressed as mean +/− SEM, statistical details can be found in the figure legends and n’s refer to the number of animals for each condition. Differences were considered statistically significant at p<0.05. All variables were analyzed using the advanced analytics software package Statistica (StatSoft, Tulsa, OK, USA).

A simple linear regression analysis was performed to illustrate the relationship between variation in z-score to distance from escape hole. A two-tailed paired two-sample Student’s t-test was used to compare delta z-score 5 seconds following a correct versus incorrect investigation and meal pattern analysis and operant responding for sucrose rewards in response to treatment in the model of acute disconnection of the CA1v to LS pathway. For measures during memory probes in the HPCv lesion and CA1v-mPFC experiments, 5-day averages for meal pattern analysis in the model of chronic disconnection and c-Fos counts following administration of CNO or aCSF, differences between groups were evaluated using two-tailed independent two-sample Student’s t-tests. For CA1v to LS acute and chronic disconnection experiments, measures during spatial foraging and spatial escape task probes, the STFP paradigm, the zero maze paradigm, and the open field test, differences between groups were evaluated using one-way ANOVAs. For the variation in photometry signal across time, meal pattern analysis for the chronic disconnection model and all measures of food intake, body weight, and errors/latency during spatial foraging and spatial escape task training, differences between groups were evaluated using two-way repeated measures ANOVAs (treatment x time). Significant ANOVAs were analyzed with a Fisher’s LSD posthoc test where appropriate. For all memory probes, as well as the STFP, one-sample t-tests were conducted to determine whether performance was significantly different from chance set at 0.1667. Outliers were identified as being more extreme than the median +/− 1.5 * interquartile range. For all experiments, assumptions of normality, homogeneity of variance (HOV), and independence were met where required. motivation, such that each hidden tunnel contained five 45mg sucrose pellets (cat. no. F0023, Bio-Serv, Flemington, NJ).

In the event that a rat did not find the correct hole during the 3min training session, the experimenter gently guided the rat to the correct hole and were allowed to crawl in and then complete the training in the same fashion as rats that found the hole on their own. During training sucrose pellets were taped underneath all of the holes to preclude olfactory-based search strategies during training. Importantly, the memory probe test does not include any sucrose

---

**Supplementary Figure 1.**
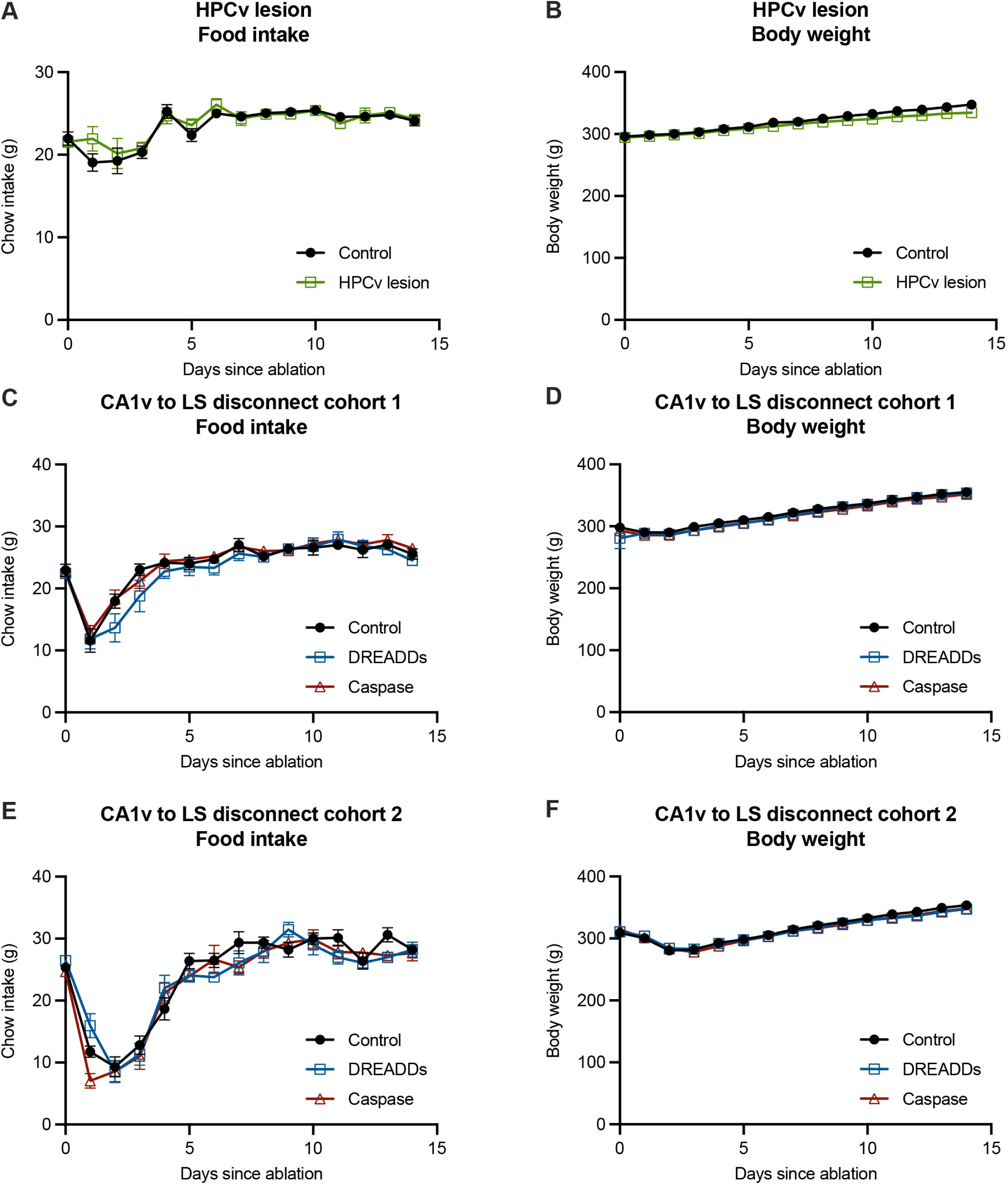
Effect of bilateral HPCv lesions and reversible and chronic disconnection of the CA1v to LS neural pathway on food intake and body weight. There were no effects on food intake (A) or body weight (B) of bilateral HPCv lesions compared to controls. There were no effects on food intake (cohort 1: C; cohort 2: E) or body weight (cohort 1: D; cohort 2: F) of reversible (DREADDs) or chronic (caspase) disconnection of the CA1v to LS neural pathway compared to controls. For graphs S2A-B (HPCv lesion), lesion n=11, control n=18. For graphs S2C-D (CA1v to LS disconnect cohort 1), DREADDs n=6, caspase n=10, control n=8. For graphs S2E-F (CA1v to LS disconnect cohort 2), DREADDs n=8, caspase n=12, control n=10. For graphs S2E-G, DREADDs vehicle n=4, DREADDs CNO n=4. For graphs S2H-I, caspase n=6, control n=2). All values expressed as mean +/- SEM.

**Supplementary Figure 2.**
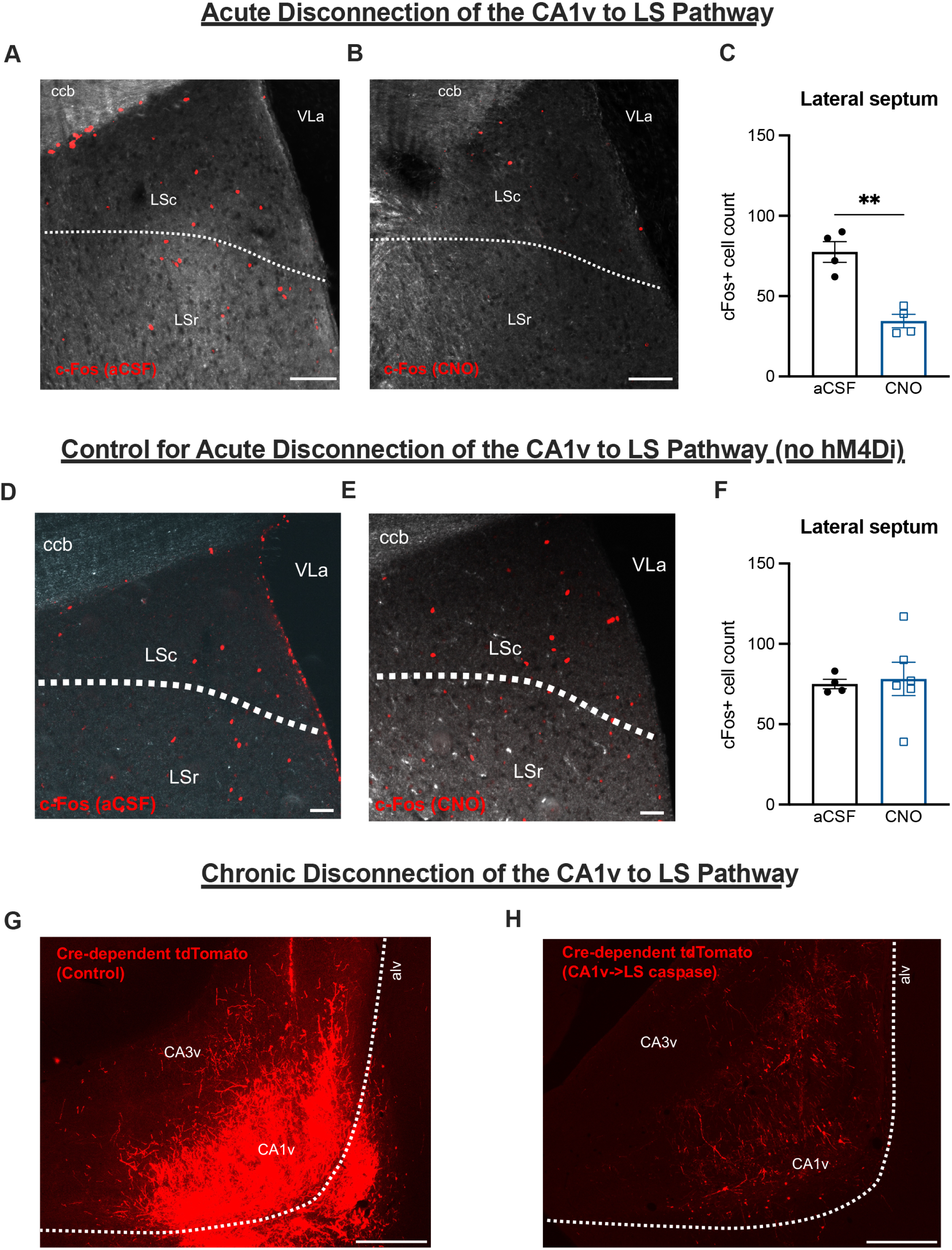
Verification of the acute and chronic approach. Histological verification of inhibitory DREADDs-induced acute silencing of LS-projecting CA1v neurons determines that the appetitive spatial memory probe induces c-Fos activation in the LS in aCSF-treated animals (A; scale bar 100μm), an outcome significantly reduced in CNO-treated animals (B; scale bar 100μm, C; p<0.01). Histological verification in animals lacking inhibitory DREADDs in LS-projecting CA1v neurons determines that the appetitive spatial memory probe induces similar c-Fos activation in the LS following ICV infusion of aCSF (D; scale bar 100μm) or CNO (E; scale bar 100μm, F). Histological verification of caspase-induced chronic silencing of the LS-projecting CA1v neurons demonstrates that cre-dependent tdTomato labeling of CA1v neurons induced by LS-origin retro-cre is robust in a control animal (G; scale bar 500μm), but substantially reduced in an animal injected with cre-dependent caspase combined with cre-dependent tdTomato due to caspase-induced cell death (H; scale bar 500μm). For graph S1C, aCSF n=4 and CNO n=4. For graph S1F, aCSF n=4 and CNO n=6. All values expressed as mean +/- SEM.

**Supplementary Figure 3.**
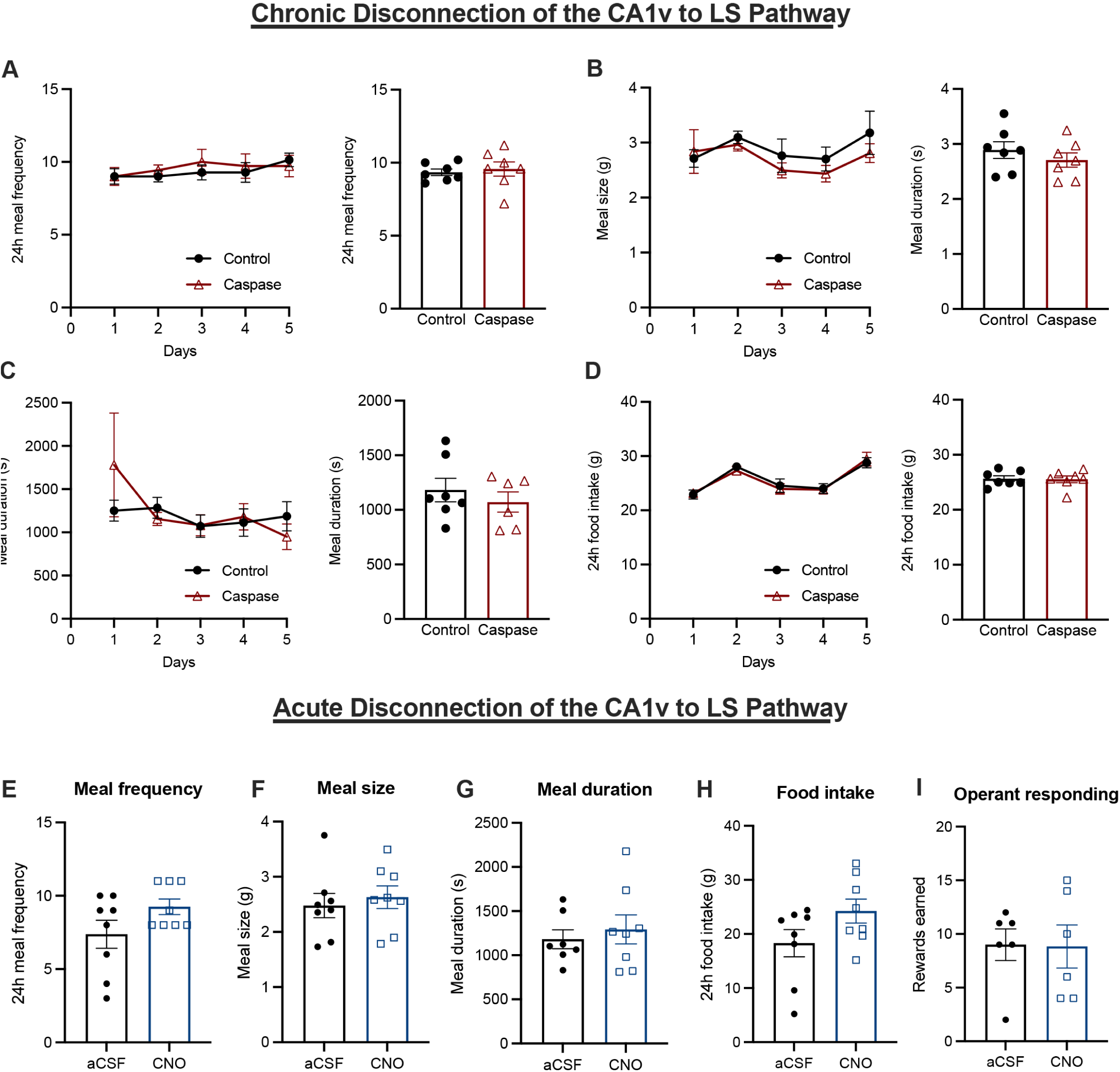
Effect of reversible and chronic disconnection of CA1v to LS neural pathway on feeding behavior and motivated operant responding for sucrose. Compared with control animals, chronic caspase-mediated disconnection of the CA1v to LS pathway did not significantly affect meal frequency (A), meal size (B), meal duration (C), or 24h cumulative food intake (D) over the course of the 5-day recording period. Acute DREADDs-mediated disconnection of the CA1v to LS pathway (via CNO infusion) resulted in non-significant trend towards increased meal frequency (E), no differences in meal size (F) or meal duration (G), a significant increase in 24h cumulative food intake (H) and no differences in rewards earned in a progressive ratio reinforcement procedure (I) in comparison with vehicle injection. For graphs S3A-D, caspase n=7 and control n=7. For graphs S3E-H, DREADDs n=8 (within-subjects). For graph S3I, DREADDs n=6 (within-subjects). All values expressed as mean +/- SEM.

**Supplementary Figure 4.**
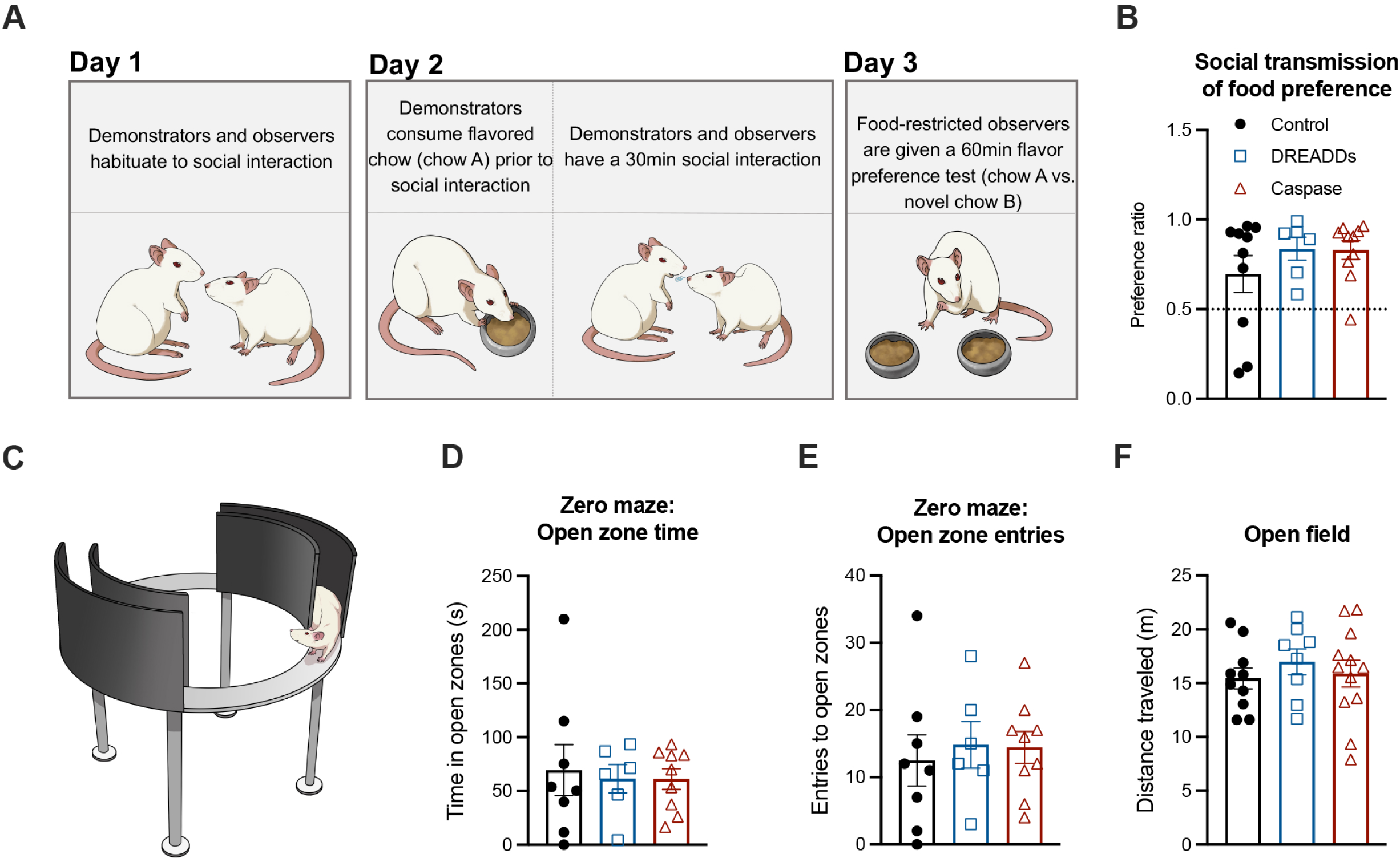
Neither reversible nor chronic CA1v to LS neural disconnection impair spatial memory for escape location, social transmission of food preference, anxiety-like behavior, or general locomotor activity levels. Diagram of the social transmission of food preference (STFP) task (A). Neither reversible nor chronic disconnection of the CA1v to LS pathway impair STFP learning compared to controls, as measured by a food preference ratio (B), with the dotted line indicating chance preference level (0.50). Diagram of the zero maze apparatus (C). Anxiety-like behavior was not influenced by reversible or chronic disconnection of the CA1v to LS pathway compared to controls, as measured by performance in the zero maze task, specifically time in open zones (D) and entries into open zones (E). Neither chronic nor reversible CA1v to LS disconnection affected open field performance compared to controls, as measured by total distance traveled (F). For graphs 3B, 3D, and 3E (CA1v to LS disconnect cohort 1), DREADDs n=6, caspase n=10, control n=8. For graph 3F (CA1v to LS disconnect cohort 2), DREADDs n=8, caspase n=12, control n=10. All values expressed as mean +/- SEM.

**Supplementary Figure 5.**
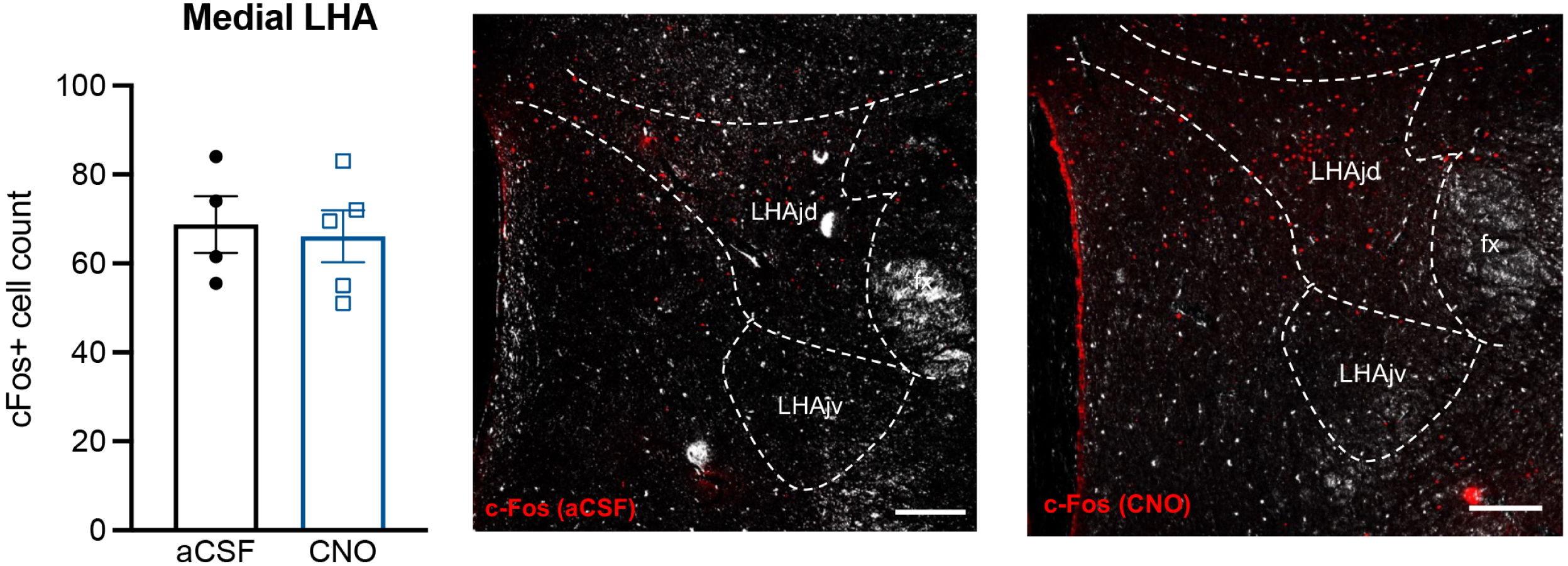
Medial LHA activation following CNO administration in the absence of DREADDs. Counts of c-Fos positive cells in the medial LHA (LHAjd and LHAjv) are unchanged with ICV administration of CNO in the absence of inhibitory DREADDs. Representative histology of c-Fos expression in the medial LHA (LHAjd and LHAjv; scale bar 200μm) following ICV infusion of aCSF (n=4) or CNO (n=6). All values expressed as mean +/- SEM.

**Supplementary Figure 6.**
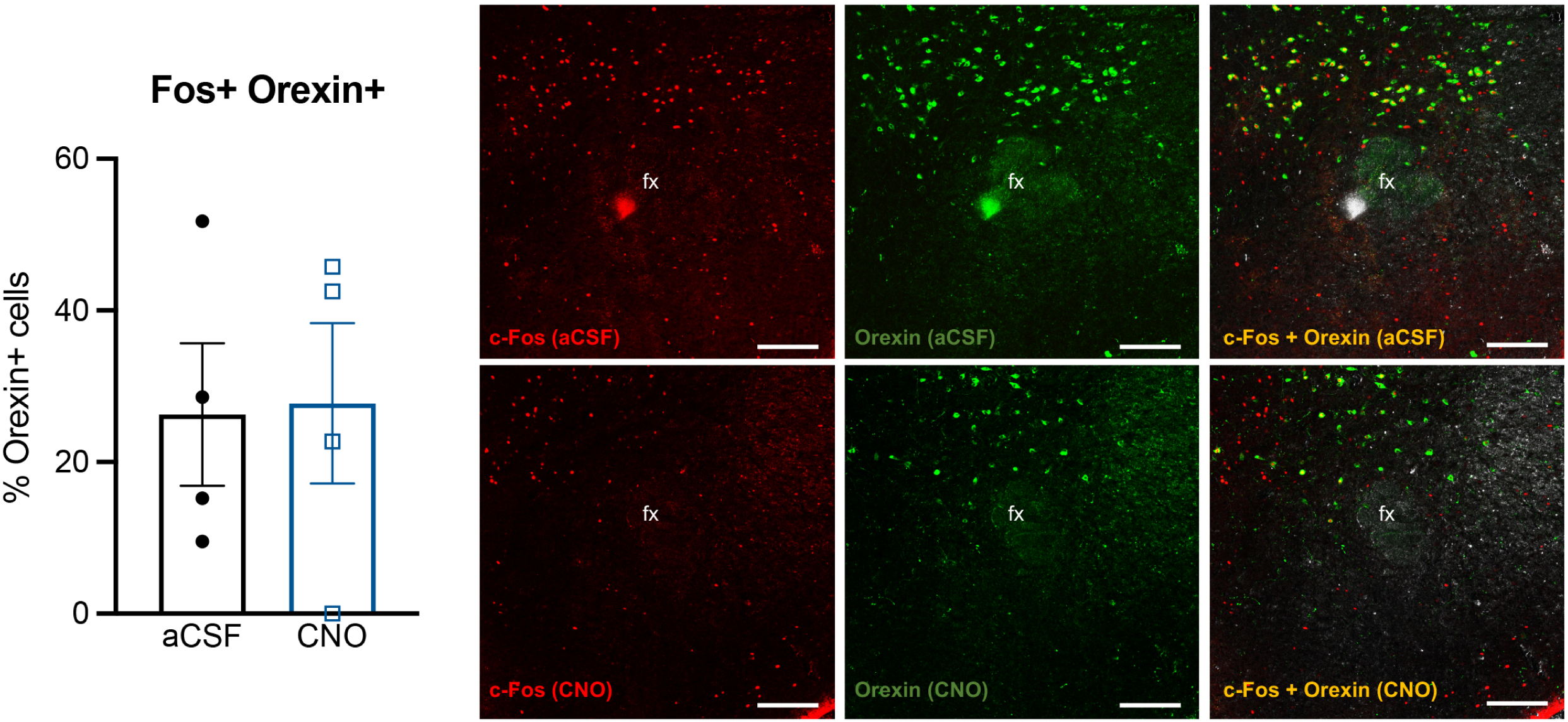
Phenotyping of medial LHA active neurons following appetitive spatial memory procedures with acute disconnection of the CA1v-LS pathway. The percentage of c-Fos positive orexin neurons in the medial LHA is unchanged with ICV administration of CNO (vs. aCSF) to acutely disconnect the CA1v-to-LS pathway. Representative histology of c-Fos and orexin expression in the medial LHA (scale bar 200μm) following ICV infusion of aCSF (n=4) or CNO (n=4). All values expressed as mean +/- SEM.

**Supplementary Table 1.**
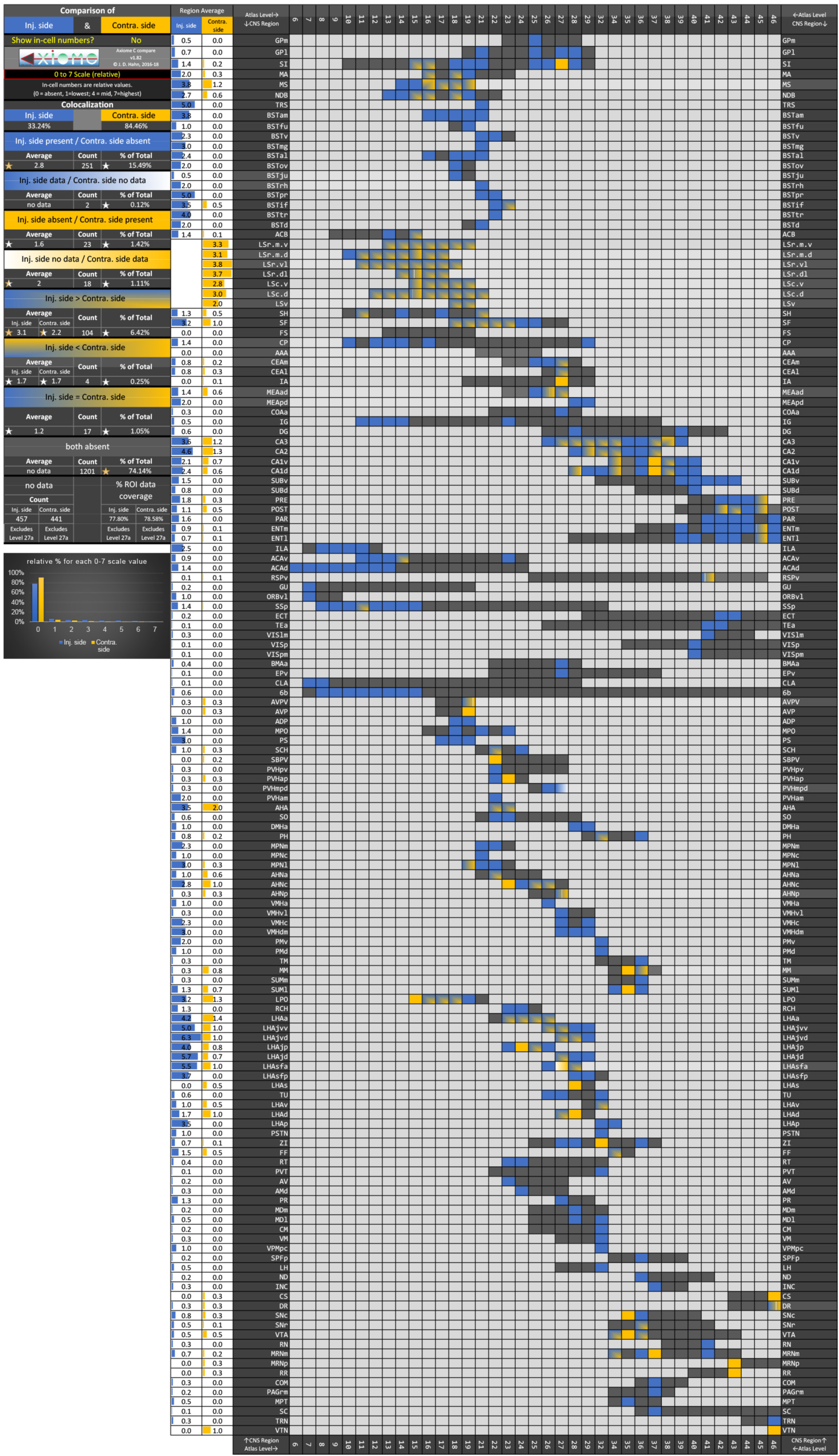
Summary table of the forebrain projection targets of LS neurons that receive input from CA1v (derived the same raw data as the brain flatmap summary diagram in Fig. 5D).

